# Distinct Cortical-Thalamic-Striatal Circuits Through the Parafascicular Nucleus

**DOI:** 10.1101/370734

**Authors:** Gil Mandelbaum, Julian Taranda, Trevor M. Haynes, Daniel R. Hochbaum, Kee Wui Huang, Minsuk Hyun, Kannan Umadevi Venkataraju, Christoph Straub, Wengang Wang, Keiramarie Robertson, Pavel Osten, Bernardo L. Sabatini

## Abstract

The thalamic parafascicular nucleus (PF), an excitatory input to the basal ganglia, is targeted with deep-brain-stimulation to alleviate a range of neuropsychiatric symptoms. Furthermore, PF lesions disrupt the execution of correct motor actions in uncertain environments. Nevertheless, the circuitry of the PF and its contribution to action selection are poorly understood. We find that, in mice, PF forms the densest subcortical projection to the striatum. This projection arises from transcriptionally- and physiologically-distinct classes of PF neurons that are also reciprocally connected with functionally-distinct cortical regions, differentially innervate striatum neurons, and are not synaptically connected in PF. Thus, mouse PF contains heterogeneous neurons that are organized into parallel and independent associative, limbic, and motor circuits. Furthermore, these subcircuits share motifs of cortical-PF-cortical and cortical-PF-striatum organization that allow each PF subregion, via its precise connectivity with cortex, to coordinate diverse inputs to striatum.

## INTRODUCTION

Selecting and generating appropriate motor-actions requires integration of limbic, associative, and sensory information in basal ganglia (BG) circuits (Macpherson et al., 2014) (Hintiryan et al., 2016), a set of phylogenetically old subcortical nuclei (Stephenson-Jones et al., 2011). The importance of these nuclei to action-selection in humans is emphasized by disorders that arise from disrupted components of the BG, such as Parkinson’s (Wichmann et al., 2011) (Kravitz et al., 2010), Huntington’s (Vonsattel et al., 1985) (Mangiarini et al., 1996), Tourette’s Disorder (Leckman et al., 2010) (Vinner et al., 2017), Obsessive-Compulsive (Saxena et al., 2001) (Ahmari et al., 2013) and addiction (Everitt and Robbins, 2005) (Hollander et al., 2010).

The BG consist of loops formed by projections from cortex (CTX) and thalamus (TH) to the input stage of the BG, the striatum (STR), which than signals via cascading inhibitory nuclei to control cortical-projecting thalamic nuclei (DeLong, 1990) (Nelson and Kreitzer, 2014) (Cowan and Powell, 1956) (Kemp and Powell, 1970) (Hunnicutt et al., 2016) (Hintiryan et al., 2016). Phylogenetically, TH and the STR pre-date the expansion of the CTX (Reiner et al., 1998) and despite the TH being approximately ten times smaller in volume than the CTX in mice (see results), it accounts for approximately a quarter of all glutamatergic synapses in the STR (Huerta-Ocampo et al., 2014). This suggests that these evolutionally conserved projections between TH and STR have a powerful functional impact on BG circuits (Minamimoto et al., 2005) (Bradfield et al., 2013a) (Kato et al., 2011) (Smith et al., 2011) (Bradfield et al., 2013b).

Within TH, the parafascicular (PF) and centromedian (CM) nuclei (two separate nuclei that at times are defined as a complex) project heavily to STR (Smith and Parent, 1986) (Berendse and Groenewegen, 1990) (Wall et al., 2013), unlike typical thalamic nuclei that primarily interact with CTX (Sherman and Guillery, 2013). In humans, targeting PF/CM for deep brain stimulation (DBS) has been successful in alleviating a range of symptoms in individuals with BG-related disorders (Testini et al., 2016) (Savica et al., 2012) (Peppe et al., 2008) (Jouve et al., 2010) (Parker et al., 2016) (Picillo et al., 2017). Furthermore, PF/CM are unique in that they degenerate early in Parkinson’s, unlike other thalamic nuclei that maintain their integrity throughout disease progression (Henderson et al., 2000a). However, the PF is omitted from the majority of functional models of the BG in both primate and rodent literature (Penney and A. B. Young, 1983) (DeLong, 1990) (Nelson and Kreitzer, 2014), or grouped together with other thalamostriatal projections, despite evidence that the anatomy and function of PF➔STR projections is specialized (Ellender et al., 2013) (Alloway et al., 2014).

In primates, the projections from PF/CM to STR have been proposed to be anatomically organized into multiple functionally distinct output channels (Steiner and Tseng, 2016) (Sadikot and Rymar, 2009), a conclusion that is broadly in agreement with anatomical findings from cats and rats (Giménez-Amaya et al., 2000) (Jones, 2007). However, understanding the polysynaptic nature of circuits across connected regions in genetically-intractable species is challenging. Therefore, although it is known that subregions of PF/CM project to different regions in STR, it has not been possible to link these specialized projections to cell classes within PF/CM or to understand their relationship to the many cortical regions that project to intralaminar (ILM) TH and STR. Thus, it is unknown if cortical-PF-STR and cortical-PF-cortical circuits are organized into conserved motifs. Furthermore, because of the limitations of genetic manipulations and *ex-vivo* electrophysiology analysis in these species, little is known about the cellular composition and micro-circuitry of PF/CM, the neurons that comprise its input and output channels, and synapses by which PF modulates STR activity. Conversely, in rodents, the lack of clear histological demarcations within PF and between PF and CM as well as the small size and close packing of TH nuclei, has led researchers to treat the PF in genetically-tractable species such as mice as anatomically uniform and cellularly homogenous (Parker et al., 2016) (Kato et al., 2011) (Aceves Buendia et al., 2017) (Assous et al., 2017) (Choi et al., 2018).

Here we combine anatomical analysis of the circuitry linking cortex, PF and STR, with transcriptional and electrophysiological analyses of PF neurons and their synapses, to deconstruct the mouse PF. Using quantitative whole-brain anatomical approaches, we reveal PF to be the densest sub-cortical input to the STR out of hundreds of brain structures (as annotated in the Allen Institute Common Coordinate Framework). Anatomical, single-cell transcriptional, and physiological analyses reveal distinct neuronal populations in PF that form topographically organized projections to the STR. Based on these results, we target each PF subpopulation to map their cortical inputs and outputs. We find that PF cell classes are targeted by layer 5 of limbic, associate or sensory-motor regions of CTX while also forming topographically organized projections to CTX. Furthermore, these PF cell classes are not synaptically interconnected within PF and differentially innervate striatal neurons, thus forming functionally-distinct and parallel signaling channels.

Our circuit analyses reveal that PF subregions and neuron classes influence distinct regions of STR through independent and parallel channels that carry information principally from limbic, associative, or sensorimotor regions. These channels are organized such that an area of STR receives input from regions of CTX and PF that are themselves interconnected via reciprocal projections. Based on this organization, we propose that PF circuits facilitate and dynamically shape the output of connected and behaviorally relevant striatal regions to mediate correct action selection in the ongoing sensorimotor context.

## RESULTS

### Quantification of the distribution of inputs to striatum across the brain

The input nucleus of the BG, the STR, receives inputs from many parts of the brain, including CTX and TH (Steiner and Tseng, 2016). These inputs were previously mapped using retrograde tracing and manual cell counting to quantify inputs to STR from a few dozen brain regions (Wall et al., 2013). Alternatively, anterograde tracing was combined with image analysis to identify functionally distinct regions in the STR defined by the combination of inputs that they receive from CTX and TH (Hunnicutt et al., 2016) (Hintiryan et al., 2016).

We utilized automated image acquisition and analysis to map the distribution of putative STR-projecting neurons across the whole brain (Figure 1 and Movie 1). We injected 4 locations in the STR of 7 C57BL6/N wild type (WT) mice with a non-pseudotyped rabies virus encoding nuclear localized GFP (RV-nGFP) (Figure 1A). The 3D whole brain volume was subsequently imaged, reconstructed, and aligned to the Allen Brain Atlas (ABA) for analysis (Figure 1A). As our data is aligned to the ABA, here we use their brain-structure hierarchy and abbreviations defined by the Allen Institute (link: <u>ABA interactive atlas</u> <u>viewer</u>) with the exception of brain-stem which is switched here with sub-cortical (sub-CTX; see Table 1 for all brain structure abbreviations used in figures).

**Figure 1:**
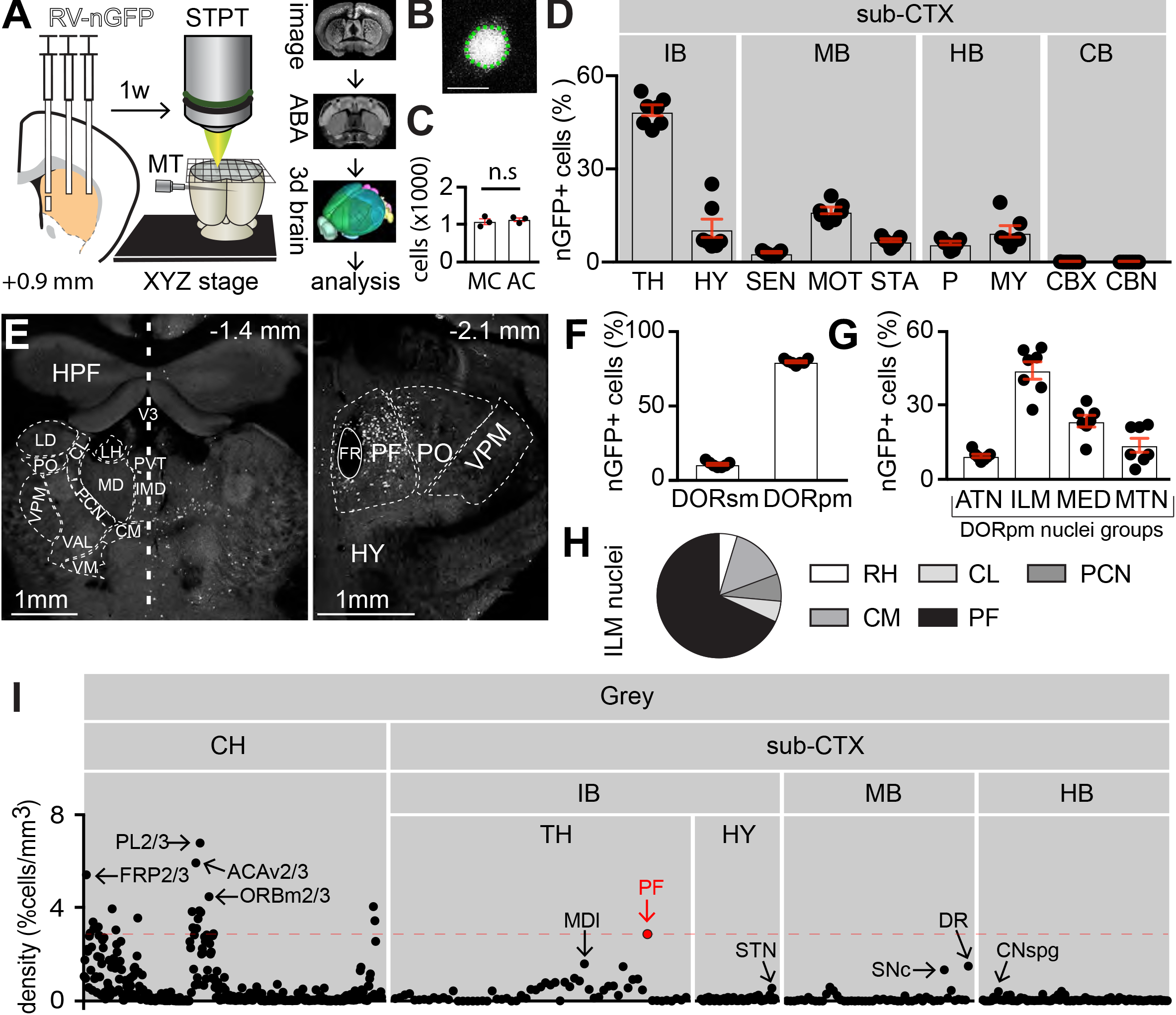
Serial two photon tomography defines PF as the main sub-cortical input to STR. **A**, *left*, Schematic of the experimental design showing a coronal section at +0.9 mm from a WT mouse with 4 injections of RV-nGFP in the STR (region of injection is highlighted in orange in this and in all subsequent figures). *middle*, Schematic of the STPT system, which automatically slices and images the whole brain using a microtome (MT) built into a 2-photon laser-scanning fluorescence microscope. *right*, Image of a brain slices obtained approximately 1 week after virus injection that was aligned to the ABA and 3D reconstructed for further analysis. **B**, STPT image of the nucleus of a cell infected with RV-nGFP (white). The boarder of the nGFP and the tissue are marked with a dashed line (green) to highlight the signal to noise ratio obtained with RV-nGFP despite the dense labeling neurons upstream of STR. **C**, Number of cells detected in PF by manual (MC) and automated (AC) counting (n=3 mice). P=0.5; Wilcoxon test. Red error bars in this and subsequent panels indicate ±SEM and black bar indicates the mean. In this panel and D, F and G each black dot indicates data from one mouse. **D**, Percent RV-nGFP+ cells in subcortical (sub-CTX) regions from the experiment shown in panel (A). Each dot shows the percentage of cells in the indicated brain region measured in one mouse (n=60,857/7; cells/mice). See Table 1 for full list of abbreviations. **E**, Coronal sections of the TH at -1.4 mm (*left*) and -2.1 mm (*right*) showing RV-nGFP infected cells (white). On the left, boundaries of TH nuclei are shown with thin dashed lines contralateral to the injection site in STR to not obscure the nGFP signal in cells across the many nuclei. The thick dash line represents the midline. See Table 2 for full data set. **F**, Percentage of total TH RV-nGFP+ cells found in sensory-motor cortex related (DORsm) or poly-modal association cortex related (DORpm) regions of TH (n=26,490/7; cells/mice). **G**, Percentage of total DORpm RV-nGFP+ cells found in 4 of the DORpm nuclei groups (n=23,191/7; cells/mice). **H**, Pie chart of distribution of RV-nGFP+ cells across the nuclei of the intralaminar (ILM) TH (n=10,019/7; cells/mice). PF is labeled in black. **I**, Relative cell density (defined by percent of total RV-nGFP+ cells in each brain region divided by the region’s volume) for 706 regions. Each black circle represents the mean cell density across animals (n=7). Regions with high densities of RV-nGFP+ cells are highlighted with arrows. PF is highlighted in red and has the highest density of putative projection neurons to STR in the sub-CTX as seen by the red dashed line that marks PF density (n=668,890/7; cells/mice). Hierarchical organization of brain regions and abbreviations are based on the Allen Brain Atlas Interactive Atlas Viewer (link: <u>ABA Interactive Atlas Viewer</u>). Abbreviations look up table for this Figure and all others is in Table 1 and the full dataset of experiment shown in panel A is in Table 2. Also see related Figure S1, and Movie 1.

The high signal-to-noise ratio (SNR) of somatic GFP signal versus the neuropil signal was exploited to automatically count RV-nGFP+ cells (Figure 1B), permitting an unbiased estimate of putative inputs to STR across the whole brain and from hundreds of structures (Movie 1). The coefficients of variation (CV) of the volumes of 8 brain-regions of interest across mice were less than 4% (Figure S1A-C) allowing pooling of data across brains. The false positive rate (FPR) for automated detection of RV-nGFP+ cells was estimated from cell counts in the STR and PF contralateral to the injection site as there is no PF➔STR or STR➔STR connectivity across hemispheres. This yielded an estimate of <1% FPR (ipsilateral counts: STR=23396±2332 cells, PF=6797±81 and contralateral counts: STR=229±42, PF=51±1; n=7 mice; Figure S1D). Furthermore, in a subset of animals. labeled PF neurons were counted manually, yielding numbers very similar to the automated measurements (manual: 3219±80 cells; automated: 3375±48; n=3 mice; P=0.5; Figure 1C).

To investigate the distribution of inputs to STR in the sub-CTX main hierarchical divisions, the percentage of the total RV-nGFP+ cells located in each group was calculated. TH had the highest percent of nGFP-labeled cells with the motor region of the midbrain (MBmot) being second (TH=48±1% of cells; MBmot=16±1%; see Table 2 for full dataset) compared to the other 7 sub-CTX groups which together had 35% of putative sub-CTX input to STR (Figure 1D; see Table 1 for full list of abbreviations for this figure and all others). In TH, the majority of nGFP+ cells were in the poly-modal association cortex-related region (DORpm) and not the sensory-motor cortex-related region (DORsm) (DORpm=79±0.6%; DORsm=10±0%; Figure 1E-F). Among DORsm nuclei, the ventral anterior lateral complex had the most cells (S1E), similar to previous observations in the squirrel monkey (Smith and Parent, 1986).

In DORpm, the intralaminar nuclei group (ILM) had the majority of putative STR inputs (43±3%; Figure 1G) with PF having the highest percent of nGFP+ cells compared to all other ILM nuclei (68±1% of cells; Figure 1H). Lastly, the density of putative inputs to STR from 706 ABA-defined sub-structures was calculated (defined as % of total cells in a given region divided by its volume) and showed that PF had the densest sub-CTX input to STR, highlighting its potential to exert powerful control of the BG circuits (Figure 1H and Table 2).

### The parafascicular-striatal projections are topographically organized in mice

In cats, primates, and humans, the PF/CM complex is separated into histologically dissimilar PF and CM nuclei which are topographically organized into multiple output channels that target distinct regions of the STR (Jones, 2007) (Steiner and Tseng, 2016). To address whether the PF/CM➔STR topography found in larger species exists in mice, WT mice were injected with 3 variants of the retrograde tracer cholera toxin subunit B (CTB) (Conte et al., 2009) into four STR locations (Figure 2A). Topographically organized projections were observed in the anterior part of PF (coronal section -2.0mm defined by the ABA as the anterior posterior boarder of PF with the medial dorsal nucleus) between medial PF (mPF) and medial STR (mSTR); central PF (cPF) and dorsal medial STR (dmSTR); and lateral PF (lPF) and dorsal lateral STR (dlSTR) (Figure 2B-D). This topography was maintained at coronal section -2.1mm (Figure 2E-G) and -2.2mm (Figure 2H-J) across animals but becomes less distinct at coronal section -2.3mm, corresponding to the most posterior part of both PF (and the TH) (Figure 2K-M). The topography of projections from cPF➔dmSTR was also clearly seen in cleared brains (Movie 2) (Chung et al., 2013). Lastly, the ventral part of mPF (v-mPF) projects to the nucleus accumbens (ACB) (Figure S2A-B). Thus, despite its small size, and no clear boundary to define CM and PF, PF in mice contains distinct and topographically organized PF➔STR projections that share similar organizational features with larger species (Jones, 2007) (Giménez-Amaya et al., 2000).

**Figure 2:**
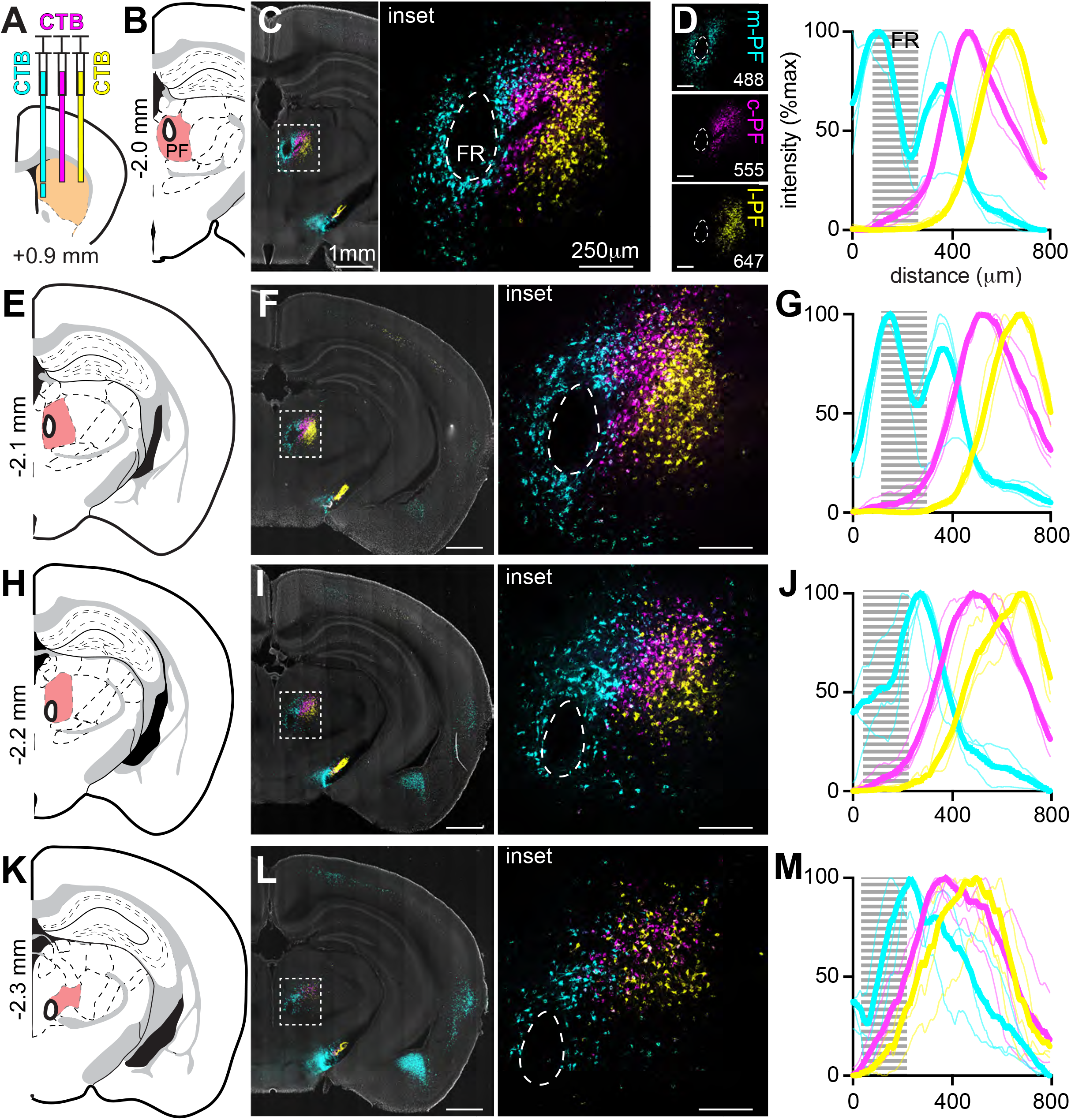
PF➔STR projections are topographically organized in mice. **A**, Schematic of the experimental design showing a coronal section at +0.9 mm from a WT mouse with 4 injections of 3 CTB variants (cyan, magenta, and yellow) in the STR. The region of injection is highlighted in orange. **B**, Coronal section from the ABA at -2.0mm with PF highlighted in red and the fasciculus retroflexus (FR) circled with a thick black line inside of PF. The FR was used as a landmark in PF to align images for summary data shown in panels D, G, J, and M. **C**, Example image of a coronal section at -2.0mm (*left*) from the experiment shown in A with the inset indicating the region surrounding the PF enlarged on the *right*. The distributions of CTB conjugated with different fluorophores are largely not overlapping, highlighting the PF➔STR topographical organization. **D***, left*, Confocal image of PF tissue excited with 488, 594, and 647 nm wavelengths (*top* to *bottom*) highlighting the topographical organization of the PF-STR projections. *right*, Quantification of fluorescence intensity for each imaging channel at coronal section - 2.0mm along the medial-lateral axis. Thin lines represent peak-normalized data from individual animals and the thick lines show the means for each channel. The dashed grey region represents the FR location. Scale bar = 250 μm (n=3 mice). **E-G**, Atlas schematics, example images, and quantifications as in B-D for coronal sections -2.1 (E), -2.2 (F), and -2.3 (G) mm. The example images are from the same mouse shown in panels (C-D). Also see related Figure S2, and Movie 2.

### Characterization of PF neuron types using single cell sequencing and electrophysiological interrogation

To examine the neuronal heterogeneity in PF we used a droplet-based single cell RNA sequencing technique (inDrops) (Klein et al., 2015) (Hrvatin et al., 2018). This technique allowed us to determine whether cells in PFs sub-divisions, defined by the PF➔STR projections topography, are transcriptionally distinct. PF and its surrounding areas were manually dissected from acute coronal brain slices and a cell suspension was formed by tissue dissociation (Figure 3A). Analysis of transcriptomes of 10,471 cells from 8 mice, revealed 7 main cell classes with distinct transcriptional profiles (Figure 3B). The neuronal cell-type class (enriched for *Snap25*, *Syn1*) contained 992 cells and expressed markers for glutamatergic (e.g. *Slc17a6*), but not GABAergic neurotransmission (e.g. *Slc32a1*). Further sub-clustering of the neuronal cell-type class revealed 3 neuronal subclasses with cell-type enriched gene expression (see Methods) (Figure 3C). Examination of the expression patterns of the genes enriched in each subcluster in the ABA *in situ* hybridization (ISH) database (<u>Link:</u> <u>ABA ISH</u>) (Lein et al., 2007) revealed genes both inside and outside of PF. Genes whose expression is elevated in Cluster 1, including *Tnnt1,* are expressed outside of PF, primarily in posterior complex and the ventral posteromedial nucleus of TH (Figure 3D; Table 3) (Phillips et al., 2018). Genes defining cluster 2, including *Fxyd6*, were expressed in mPF, but also ventral and dorsal to the PF (Figure 3D; Table 4). Thus cluster 1 and 2 are enriched for genes and represent cell groups that, within the dissection area, are not unique to the PF. No further analysis of these clusters was carried out.

**Figure 3:**
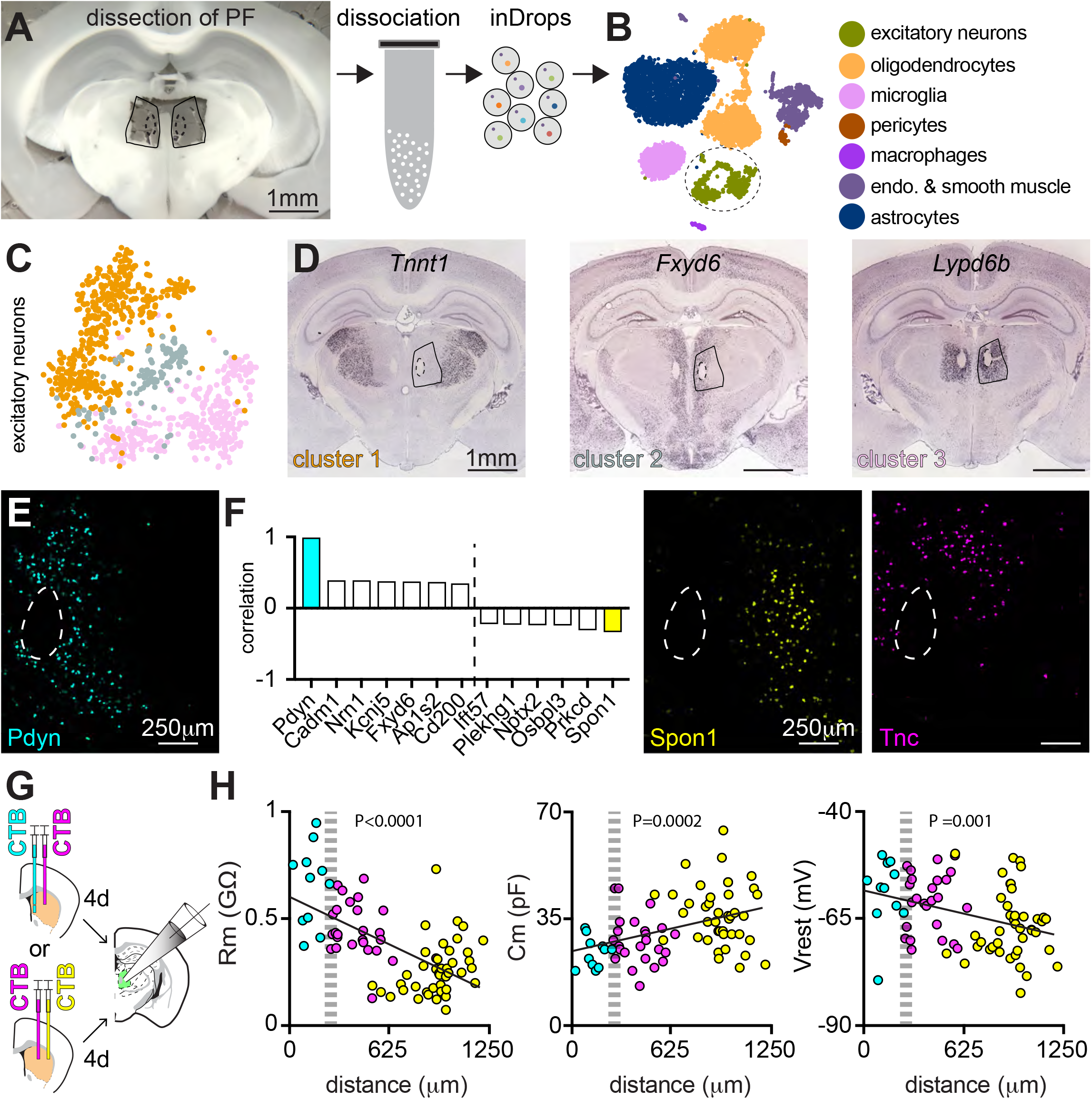
Transcriptional and electrophysiological characterization of PF neurons. **A**, *left*, Images of an acute coronal slice after microdissection of PF. *right*, Cell suspensions were formed from the dissected tissue and run through the *inDrops* platform to reveal transcriptomes of thousands of cells from PF. **B**, t-SNE plot showing the main identified cell types highlighted by the different colors (n=10471/8 cells/mice). The excitatory neurons are marked by the dashed oval delineates the neuronal cluster. **C**, t-SNE plot of excitatory glutamatergic neurons (*Slc17a6*-expressing) with the 3 subclusters indicated by different colors (n=992/8; cells/mice). **D**, Example *in situ hybridization* (ISH) from the ABA with one gene representing each neuronal cluster. *Tnnt1* (Cluster 1; also see Table 3) is expressed in thalamic neurons outside of PF. *Fxyd6* (Cluster 2; also see Table 4) is expressed in mPF neurons and ventral dorsal to PF whereas *Lypd6b* (Cluster 3; also see Table 5) is a general PF neuronal marker. **E**, *Pdyn* is expressed in mPF neurons as shown by *ISH*. **F**, Multiple genes show significant correlation or anti-correlation with *Pdyn* expression on a cell-by-cell basis (*left*). This analysis reveals *Spon1* as being anti-correlated (yellow) with *Pdyn* and expressed in lPF whereas other genes, such as *Tnc,* are markers for cPF, as confirmed by ISH (*middle* and *right,* respectively; see Table 6). **G**, Schematic of a coronal section at +0.9 mm from a WT mouse depicting 2 experimental configurations to use CTB to label neurons from mPF and cPF (*top*) or from cPF and lPF (*bottom*) that project to STR. 4 days after injections, whole-cell recordings were made in an acute brain slices of PF (highlighted in green). **H**, Summary of intrinsic neuronal parameters (Membrane resistance (Rm), Membrane capacitance (Cm), and resting membrane voltage (Vrest)) as a function of the location and labeling with CTB of the neuron in the medial-lateral axis of the PF. The location of FR is indicated by the gray dashed area. Also see related Figure S3, and Tables 3-6.

Genes enriched in cluster 3, such as *Lypd6b*, showed specific expression in PF (Figure 3D; Table 5), including all of its subdivisions. Nevertheless, within cluster 3 genes revealed differential expression along the medial-lateral aspect of PF indicating a heterogeneous neuronal population. For example, *Prodynorphin* (*Pdyn*), a marker for the direct striatal projection neurons (dSPNs) in STR (Gerfen and W. S. Young, 1988) was expressed in 117 cells, with a mean 8-fold increase in its expression compared to neuronal clusters 1 and 2. ISH of *Pdyn* mapped the expression specifically to mPF (Figure 3E). Furthermore, analysis of gene-gene expression correlation across all cells in cluster 3 revealed those correlated with *Pdyn* expression also mapped to mPF (Figure 3F; S3A-C; Table 6). Conversely, genes whose expression was anti-correlated with that of *Pdyn,* mapped to cPF and lPF (Figure 3F; S3D-F; Table 6). These results indicate that the anatomically-defined subdomains that comprise PF map onto transcriptionally distinct subclasses of neurons.

*In vivo* recordings in primates have revealed different kinetics of activation of PF and CM neurons (Matsumoto et al., 2001). Therefore, we examined if the intrinsic electrophysiological properties of neurons projecting to the STR differ along the mediolateral aspect of the mouse PF. Whole-cell recordings in current-clamp were obtained from CTB labeled PF➔STR neurons in acute brain slices from mice injected with two different CTB colors into mSTR and dmSTR or dmSTR and dlSTR (Figure 3G). Consistent with our hypothesis, the membrane resistance, capacitance, and resting potential varied across the PF with higher input resistance, lower capacitance, and higher resting potential neurons found in the medial relative to the lateral aspects of the PF (Figure 3H). Thus, the same synaptic current would result in a larger synaptic potential in the higher input resistance mPF neurons. Coupled with the more depolarized resting potential, this suggests that mPF neurons are likely more excitable than those in the lPF consistent with *in vivo* recordings in the primate (Matsumoto et al., 2001).

### *Prodynorphin* expressing cells are located in the mPF and synaptically target the matrix of STR

The restricted expression pattern of *Pdyn* in PF and the existence of a well-characterized knock-in mouse that expresses Cre recombinase from the *Pdyn* allele *Pdyn-IRES-Cre* mouse (Krashes et al., 2014) potentially permits specific manipulation of mPF circuitry. Indeed, injection of Cre-dependent AAV (creOn-GFP) in PF of the adult *Pdyn-IRES-Cre* mice (Figure 4A) resulted in GFP expression that was restricted to mPF (Figure 4B-C, S4A-D), including in the anterior-posterior (An-Po) axis of TH (% of cells: An to PF = 6%; PF = 90%; Po to PF = 3%; n=2 mice; Figure 4D). Additionally, ISH for *Pdyn* and *Slc17a6* (vglut2), showed that the *Pdyn+* cells are glutamatergic (% of cells *Pdyn*+/*Slc17a6*+ = 98%; n=125/5/2; cells/slices/mice; Figure S4E-F). mPF *Pdyn+* cells target the medial band of STR (mSTR), from dorsal STR to the ACB, and, as shown by optogenetic activation of synaptic currents, densely innervate STR neurons (Figure 4E-H). Indeed, optogenetic stimulation of the Chr2-expressing *Pdyn*+ axons evoked excitatory post synaptic currents (EPSCs) in SPNs in mSTR but not dmSTR or dlSTR SPNs (EPSCs in mSTR neurons: 24/33; dmSTR: 0/7; dlSTR: 0/7. n=4; mice; Figure 4G-H), verifying that the *Pdyn+* cells in mPF target a specific region of the STR and ACB. The fluorophore-labeled axons of mPF *Pdyn+* neurons were not uniform within the mSTR, suggesting potential differential targeting of patch (striosome) and matrix compartments (Herkenham and Pert, 1981). Indeed, we found little overlap between GFP-labeled mPF axons in STR and regions expressing mu-opioid receptors (MOR), a marker of patches (Pert et al., 1976) (Figure 4I). Fluorescence inside of each patch compared to that of a “peri-patch” shape (100 μm wide) surrounding each patch (Figure 4I) was consistently higher for the MOR channel (log fluorescence MOR=0.14±0.01) and lower for the GFP channel (log fluorescence GFP=-0.13±0.00; n=38/9/3; patches, slices, animals), consistent with *Pdyn*+ mPF fibers avoiding the STR MOR-rich compartments.

**Figure 4:**
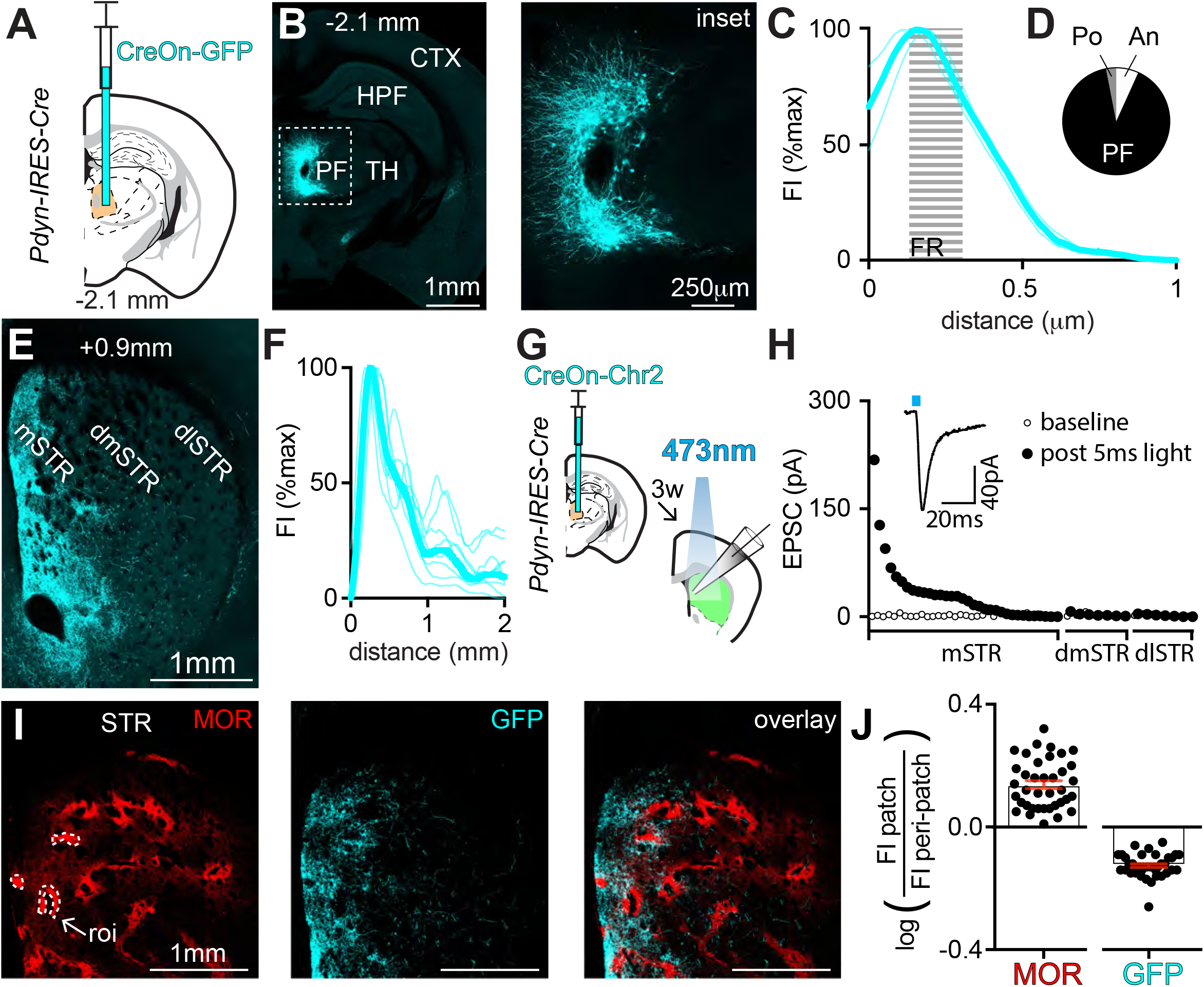
*Prodynorphin* expressing cells are located in the mPF and synaptically target the matrix of STR. **A**, Schematic of a coronal section at -2.1 mm from a *Pdyn-IRES-cre* mouse depicting an injection of creOn-gfp (cyan) AAV into the PF (injection site in PF highlighted in orange). **B**, *left,* Example coronal section at -2.1 mm of TH showing expression of GFP (cyan), indicating restricted expression of GFP in mPF. The inset is enlarged on the *right* and shows medially projecting neuronal processes from the GFP-expressing neurons. **C**, Quantification of fluorescence intensity (Fl) in PF at coronal section -2.1 mm from images such as in panel B. Thin lines represents data from individual animals and the thick lines represent the mean. The dashed grey line represents the FR location relative along the medial-lateral axis (n=3; mice). **D**, Percent of GFP+ cells anterior (An) and posterior (Po) to PF and in PF from the experiment shown in A. (n=1670/2; cells/mice). **E**, Image of a coronal section highlighting the STR at +0.9 mm from a mouse manipulated as in panel A. Dorsal STR is separated into sub-regions: Expression of GFP-expressing *Pdyn+* axons (cyan) from PF are seen in the medial STR (mSTR) but not dorsal-medial STR (dmSTR) and dorsal-lateral STR (dlSTR). **F**, Quantification of fluroescence in the STR of axons from *Pdyn*+ cells PF at coronal sections between +0.6 mm and +1.2 mm. Thin lines represents data from individual animals and the thick line represent the mean (n=9/3; slices/mice). **G**, Schematic of a coronal section at -2.1mm (left) depicting injection of AAV encoding Cre-dependent channelrhodopsin (creOn-Chr2) into PF of a *Pdyn-IRES-cre* mouse. Three weeks after virus injection whole-cell recordings were obtained in STR (green) at and around coronal section +0.9 mm. **H**, EPSC amplitudes evoked by optogenetic stimulation of *Pdyn+* PF axons and measured in SPNs as a function of region in STR (mSTR, dmSTR, dlSTR). EPSCs were recorded at -70 mV. For each cell the baseline current (open circle) and EPSC following a 5ms light pulse (closed circle) are plotted. Inset shows the mean of 10 light-evoked (blue-line indicates the light pulse) EPSCs from one cell. (n=48/4; cells/mice). Within each striatal region, results are shown ranked from largest to smallest EPSC amplitude. **I**, Image of a coronal section of the STR at +0.9 mm with mu opioid receptors (MOR) immunolabeled (red, *left*) with 3 patches in the STR highlighted (white dashed lines). Axons of *Pdyn+* PF neurons expressing GFP (center) avoid the MOR-rich patches (*overlay,* right). **J**, Quantification of the distribution of fluorescence from GFP labeled PF➔mSTR axons in and around the MOR-rich patches. The log of the ratio of the mean MOR and GFP fluorescence in the patch to that in a 100 μm wide ring around the patch (peri-patch) is shown 38 patches (n=9/3 slices/mice). All data are represented as mean (bar in black) ± SEM (red). Also see related Figure S4.

### No interconnectivity between PF cell classes

The single cell transcriptional data identified only excitatory neurons within the PF, suggesting that the topographically-organized STR-projecting neurons in PF are not interconnected by GABAergic interneurons. Several lines of analysis indicate that PF➔STR projection neurons are also not interconnected by glutamatergic synapses. First, stimulation of ChR2 in *Pdyn-IRES-Cre* mPF neurons (Figure 5A) failed to elicit light-evoked EPSCs in CTB-labeled striatum-projecting neurons in cPF (cPF➔dmSTR=0/19 EPSC; n=3 mice; Figure 5C) despite the triggering suprathreshold currents in the ChR2-expressing *Pdyn* neurons (549pA±136, 9/9 cells; n=3 mice; Figure 5B-C). Cell-filling labeling of cPF➔dmSTR projection neurons with non-pseudotyped rabies virus expressing GFP (RV-GFP) (Figure 5D) showed that axons of these neurons do not overlap with CTB-labeled STR-projecting neurons in the lPF (Figure 5D). Moreover, similar experiments using non-pseudotyped rabies expressing ChR2 (RV-ChR2) injected into dmSTR combined with a CTB injection into dlSTR (Figure 5E) resulted in light-induced currents large enough to induce action potentials (APs) in cPF neurons (237pA±56, 9/10 cells; n=3 mice; Figure 5F) but failed to evoke EPSCs in CTB+ cells in lPF➔dlSTR projection neurons (lPF➔dlSTR = 0/16 EPSC; Figure 5F). In addition, RV-mediated GFP and ChR2 expression in dlSTR-projecting lPF neurons showed no overlap of axons with CTB-labeled dmSTR-projecting neurons in cPF (Figure 5G) and no light-evoked EPSCs (cPF➔dmSTR=0/13 EPSC; n=2 mice; Figure 5H) despite suprathreshold ChR2-currents in lPF neurons (663pA±155, 6/6 cells; n=2 mice; Figure 5H-I). Lastly, trans-synaptic retrograde viral labeling with pseudotyped rabies virus (p.RV-GFP) revealed no connectivity across topographical projection zones of the PF from primary infected starter cells in mPF, or lPF. However, clear labeling in PF-projecting regions such as the Substantia Nigra Reticulata (SNr) and Superior Colliculus (SC) was observed (Figure S5).

**Figure 5:**
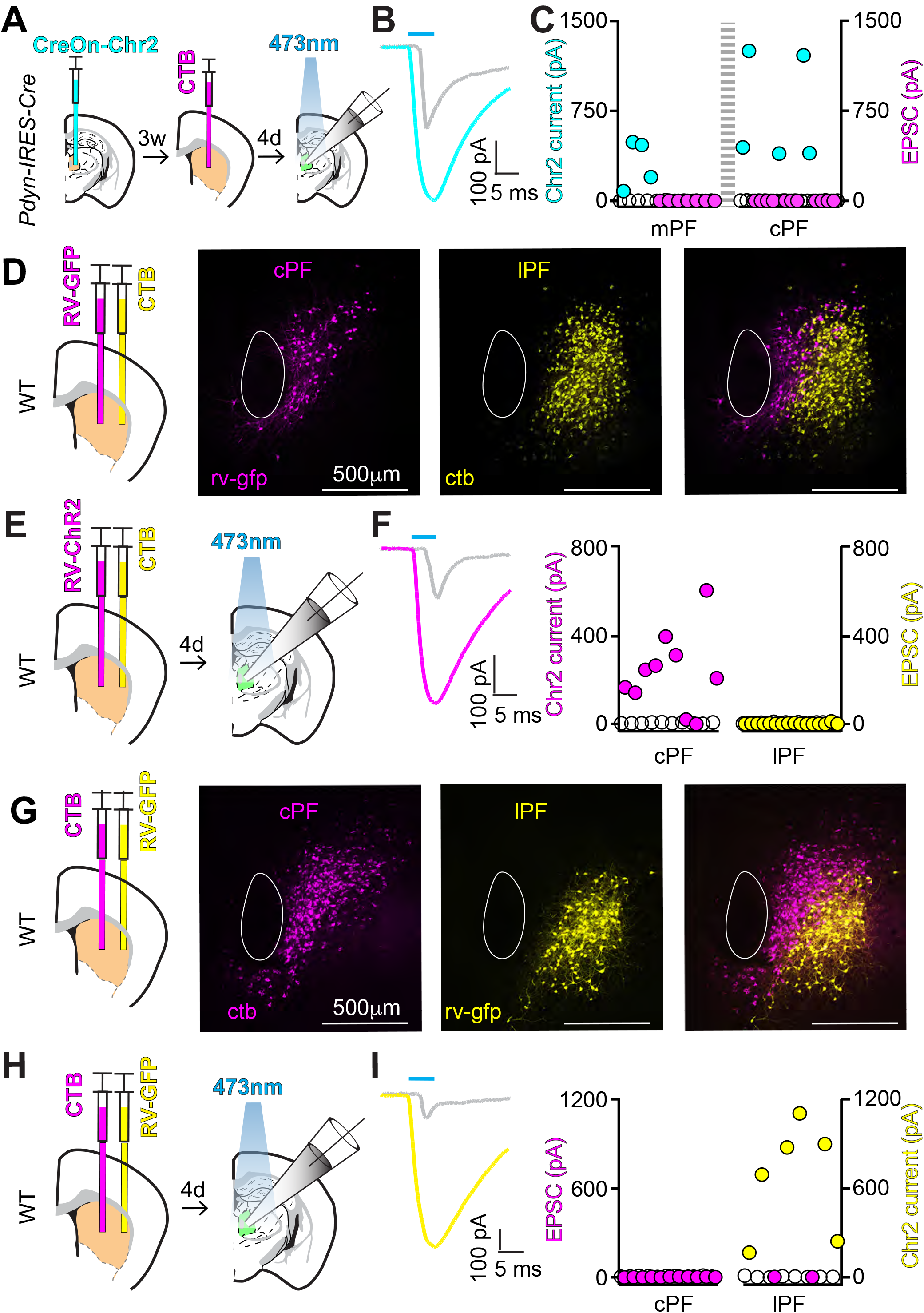
The medial, central, and lateral sub-circuits of the PF are not locally interconnected. **A**, *left*, Schematic of a coronal section at -2.1 mm depicting a viral injection of creOn-ChR2 into the PF of a *Pdyn-IRES-Cre* mouse. *center*, Coronal section at +0.9mm depicting CTB injection into dmSTR 3 weeks after the creOn-ChR2 injection. *right*, four days later acute brain slices were cut and whole-cell recordings were obtained from ChR2+ or CTB+ PF cells. **B**, ChR2-mediated excitatory currents in ChR2-expressing mPF neurons (representative example in blue) are activated concurrent with the laser pulse and display shorter latency than cortically-evoked EPSCs (representative example in gray dashed line). The cortically-evoked EPSC shown here for comparison was collected in independent the data set shown in Figure 7. **C**, EPSC (CTB+ cells) and ChR2-current amplitudes (ChR2+ cells) measured at -70 mV in mPF and cPF evoked by optogenetic stimulation of *Pdyn*-Cre+ neurons. For each cell, the baseline (white circle) and light-evoked (colored circles) currents following a 5 ms laser pulse (closed circle) are shown (n=28/3; cells/mice). The dashed grey line represents the FR and its location and circles represent the actual location of cells in the tissue relative to one another and the FR, from medial to lateral. No synaptic currents were detected in CTB+ cells (white circles for baseline and magenta circles following a 5 ms lase pulse). **D**, Experimental design showing a coronal section at +0.9 mm of a WT mouse depicting injection of RV-GFP and CTB into dmSTR and dlSTR, respectively. Images of resulting retrograde labeling in the PF (-2.1 mm) show expression of GFP (magenta) in the cPF and CTB (yellow) in the lPF. The overlay (right) shows largely not overlapped cell populations (n=3 mice, example shown from one mouse). **E**, As Panel (D) but with an injection of RV-ChR2 and followed by whole cell recordings from ChR2+ or CTB+ cells 4 days after injections. **F**, *left,* As in Panel B showing representative ChR2-mediated currents in ChR-expressing cPF neurons (magenta) compared to an example cortically-evoked EPSC *right,* As in Panel C, summary of light-evoked ChR2-mediated current (in magenta) and EPSC (yellow) current amplitudes (as in Panel C) (n=26/2; cells/mice). No synaptic currents were detected in CTB+ cells (yellow). **G**-**I**, As in Panels (D-F) but with CTB injected into dmSTR and RV-GFP or RV-ChR2 injected into dlSTR (Example images are from one of 3 representative mice. For electrophysiological analysis n=19/2 cells/mice). Also see related Figure S5.

### Subclasses of PF neurons target distinct cortical regions

Many thalamic nuclei project to and receive input from CTX forming circuits that modulate persistent cortical activity (Guo et al., 2017) (Sherman, 2016). In primates, PF neurons innervate prefrontal CTX whereas the histologically distinct CM neurons target motor and premotor areas of CTX (Parent and Parent, 2004). In rats, reconstructed cells in PF project to both the STR and several regions in CTX (Deschenes et al., 1996). However, studies of mouse PF, albeit using manipulations that could not specifically target this small thalamic nucleus, suggest that it does not project to CTX (Oh et al., 2014). To determine if PF projects to CTX in mice, and whether the subclasses of neurons that we identified in PF target distinct regions of CTX, creOn-GFP was expressed specifically in each PF cell class and automated image acquisition and analysis were used to map the distribution of GFP-labeled axons in CTX (Figure 6A). STR-projection neurons in each subregion of PF in *Pdyn-IRES-Cre* mice were targeted by a different hybrid genetic/viral intersectional strategy (Figure 6A). For mPF, Cre-dependent AAV was injected directly into PF to activate expression of GFP in *Pdyn-*expressing mPF neurons. For cPF, axon-infecting AAV encoding Flp recombinase was injected in dmSTR and AAV that expresses GFP in the presence of Flp and absence of Cre was injected into PF. This selects for neurons in cPF that target dmSTR while avoiding the nearby *Pdyn-*expressing neurons in mPF. A similar approach was used for lPF, with injection of axon-infecting Flp-encoding AAV into dlSTR. These strategies succeeded in restricting GFP somatic expression largely to the intended anterior-posterior PF region (% cells in PF for mPF targeting=83%; n=713/15/1; cPF=67%; n=1864/15/1; lPF=66%; n=2171/15/1; cells/slices/mice; Figure S6A). As expected for the PF➔STR connectivity described above, GFP+ axons were observed projecting specifically between mPF➔mSTR, cPF➔dmSTR, and lPF➔dlSTR (Figure 6B). To measure the distribution of putative PF➔CTX projections GFP+ axons were mapped to the ABA (Movie 3) and the relative axon density (RAD) was measured as fraction of all GFP+ pixels that are located in one area divided the fraction of cortical volume contained in the area. This metric gives the relative enrichment of axons in each area compared to a uniform distribution of axons within CTX.

**Figure 6:**
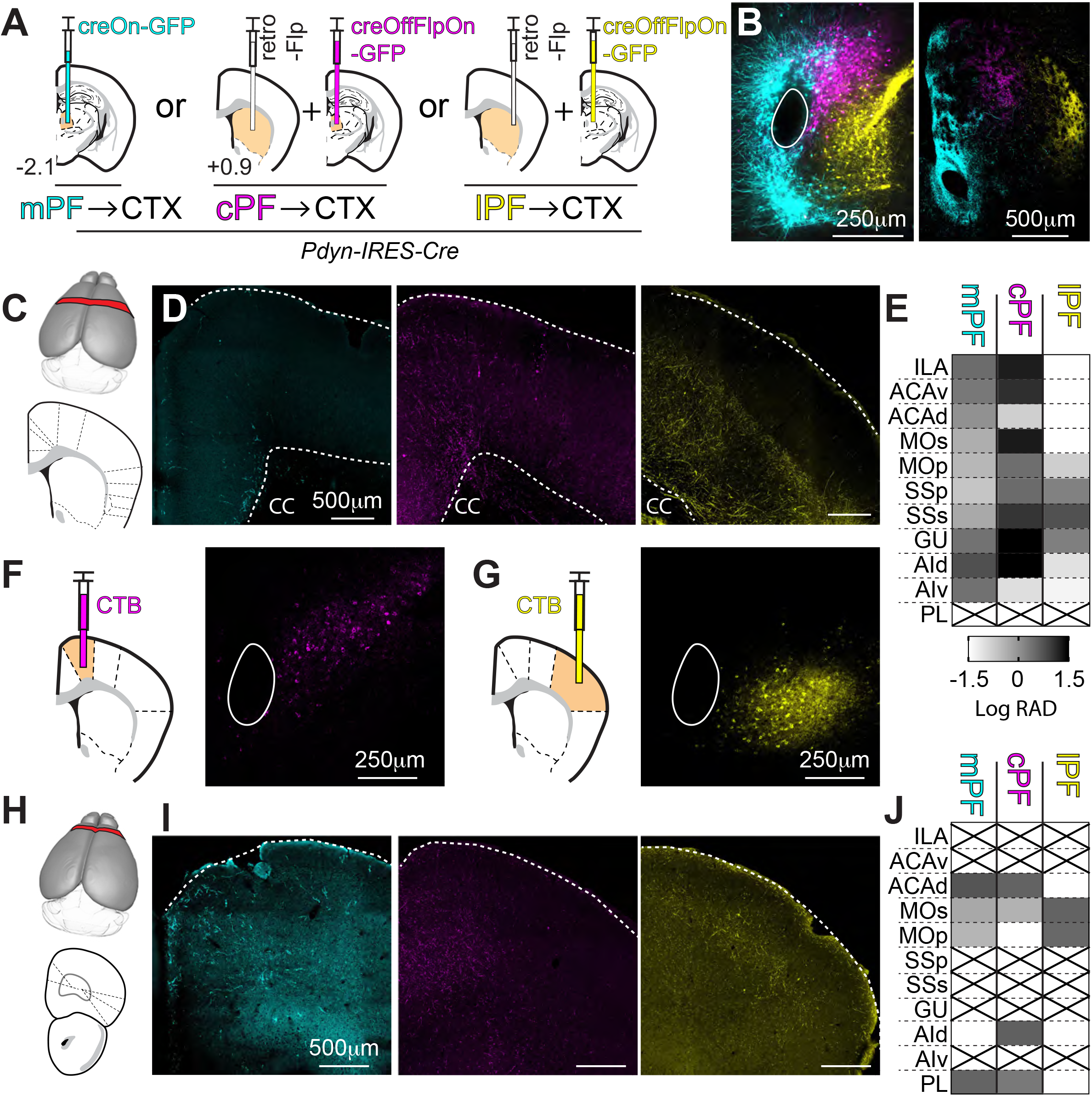
PF➔striatum projection neurons send topographically-organized outputs to CTX. **A**, Schematics of the intersectional strategies in *Pdyn-IRES-Cre* mice used to express GFP in subsets of PF➔striatum projection neurons. *left,* injection of creON-GFP (cyan) into PF results in expression of GFP in the *Pdyn* neurons which are medially located. *center,* injection of retro-Flp in dmSTR (black) and a creOff-flpOn-GFP (magenta) virus into PF ensures expression specifically in the cPF and avoids leak into *Pdyn+* neurons in mPF. *right,* injection of retro-Flp (black) in dlSTR and a creOff-flpOn-GFP virus (yellow) into PF ensuring expression of GFP specifically in the lPF. **B**, Overlay of one brain section of each of three brains targeted with the labeling strategies depicted in Panel A at coronal section -2.1 mm in PF (*left)* and at +0.9 mm in STR (*right*). **C**, *top,* For the analysis of the distribution of GFP+ axons in CTX, a region spanning from 0.6 to 1.2 mm anterior posterior was taken (red*). bottom,* Regions of interest were chosen spanning the medial lateral portion of CTX as demarcated by dashed lines. **D**, Representative coronal sections from the posterior region of CTX (0.9mm) for each of the labeling strategies (*left:* mPF➔CTX in cyan; *center:* cPF➔CTX in magenta; *right*: lPF➔CTX in yellow) highlighting the differential projections to medial, central, and lateral parts of CTX, respectively. The ventral and dorsal limits of the CTX were marked with dashes line. CC; corpus callosum. **E**, Quantification of the relative axon density (RAD) of PF axons arising for each subregion for each of 11 cortical regions. The log(RAD) per region is represented by the gray scale within caps set at ±1.5 log units. A box with an X in it indicates a cortical region not present in the analyzed slice. **F**, *left,* Experimental design showing a coronal section at +0.9 mm depicting an injection of CTB into MOs. *Right,* coronal section at -2.1mm showing CTB localized to cPF. **G**, As in Panel F but targeting SSp with CTB (*left*) resulted in labeling lPF (*right*). **H-J**, As in Panels C-E but depicting the analysis of axon distribution of an anterior section in CTX (2.5 to 3.1 mm anterior posterior). The example images shown in panels (B,D,I) are from the same mice. Also see related Figure S6 and movie 3.

We focused on 2 coronal sections of CTX (from 0.6-1.2 mm and from 2.5-3.1 mm anterior to posterior) and analyzed 11 ABA-labeled cortical subregions (Figure 6C, H). The putative output of each PF cell class was not homogenous (Figure 6D-E) in the posterior cortical section. For mPF, GFP-labeled axons were relatively enriched in the medial and lateral limbic regions of CTX (RAD from mPF to: ILA=2.8, ACAv=2.0, GU=2.41, AId=5.65 and AIv=2.42; n=1; mouse; Figure 6D-E; refer to Table 1 for abbreviations and Table 7 for full data set) and depleted in associative and sensorimotor regions (RAD from mPF to: MOs=0.4, MOp=0.3, SSp=0.2, SSs=0.4; n=1; mouse; Figure 6D-E; Table 7). cPF shared some of these medial and lateral limbic outputs with mPF but also projected heavily to MOs, GU, and AId (RAD from cPF to: MOs=18.0, GU=59.7, AId=50.2; n=1;mouse; Figure 6D-E; Table 7). In contrast, lPF projected to SSp, SSs, and GU (RAD for lPF to: SSp=1.5, SSs=5.4, GU=1.6; n=1;mouse; Figure 6D-E; Table 7) with only few axons found elsewhere in CTX. To independently verify the differential projections from PF subregions CTB was injected into posterior MOs or SSp. CTB+ cells were observed in cPF and lPF for the MOp and SSp injections, respectively (Figure 6F-G), thus confirming the results obtained with the measurements of RAD.

Similar analyses reveal that PF subregions also differentially target the more anterior section of cortex (Figure 6H). mPF projected strongly to ACAd and to PL (RAD from mPF to: ACAd=5.1, PL=3.5; n=1;mouse; Figure 6I-J; Table 7) whereas cPF shared those targets but also projected to AId (RAD from cPF to: ACAd=3.5, AId=3.9, PL=1.9; n=1;mouse; Figure 6I-J; Table 7). lPF projected to MOs and MOp of the anterior CTX (RAD from lPF to: MOs=3.5, MOp=3.1; n=1;mouse; Figure 6I-J; Table 7). Thus, striatum-projecting PF neurons differentially innervate cortical regions. mPF and cPF innervate mainly limbic structures while cPF also targets associative areas such as MOs. lPF selectively targets sensorimotor cortical areas in the posterior part of CTX and innervates MOp and MOs regions in the anterior part of CTX. This topography was generally maintained in the cortical sections between these regions (Figure S6B).

### Cortical Layer 5 projections to PF are topographically organized and form feedforward Cortex-PF-Striatum circuits

Some thalamic nuclei modulate sequential processing stages in CTX by receiving input from an upstream cortical region and projecting to its downstream cortical target, thus adding a parallel processing stage linking regions in CTX that are also themselves interconnected (Sherman, 2016) (Stroh et al., 2013) (Theyel et al., 2010). For example, in TH, the Pulvinar nucleus mediates a cortical-thalamo-cortical projection to facilitate transmission of information about attentional priorities between two visual cortical areas that are also directly connected (Saalmann et al., 2012). Since CTX is analogously upstream to the STR, we hypothesized that the cortical-thalamo-cortical circuit organization and function might also be recapitulated in cortical-thalamo-striatal circuit between CTX, PF, and STR (Saalmann, 2014).

Therefore, we examined two potential features of the circuits. First, are regions of PF and CTX reciprocally connected. Second, are regions of CTX and PF that project to the same subregion of STR themselves connected. To examine the first question – i.e. do the regions of CTX that receive input from specific subregions of PF, as defined in Figure 6, project back to those same regions of PF – we virally expressed GFP in Layer 5 projection neurons, including those that project to STR (Gerfen et al., 2013) in Tg(Rbp4-cre)KL100Gsat mice (link: <u>GENSAT resource</u>) (in short, *Rbp4cre+/-*) and labeled specific PF➔STR projection neurons by focal injection of CTB into the STR. Layer 5 neurons were targeted because they give rise the cortical outputs that participate in CTX-TH-CTX circuits described above (Sherman, 2016). Targeting MOs axons and cPF➔dmSTR cell bodies in cPF (Figure 7A) (max FI of CTB in: cPF=67±8; in rPF=8±1%; n=14/3; slices/mice; Figure 7B, S7A) revealed that MOs axons in PF preferentially overlap with CTB+ cell bodies in cPF (max fiber FI overlap in: cPF=86%±3; rPF=30%±5; n=14/3; slices /mice; Figure 7B-C) across all coronal sections of PF (S7A).

**Figure 7:**
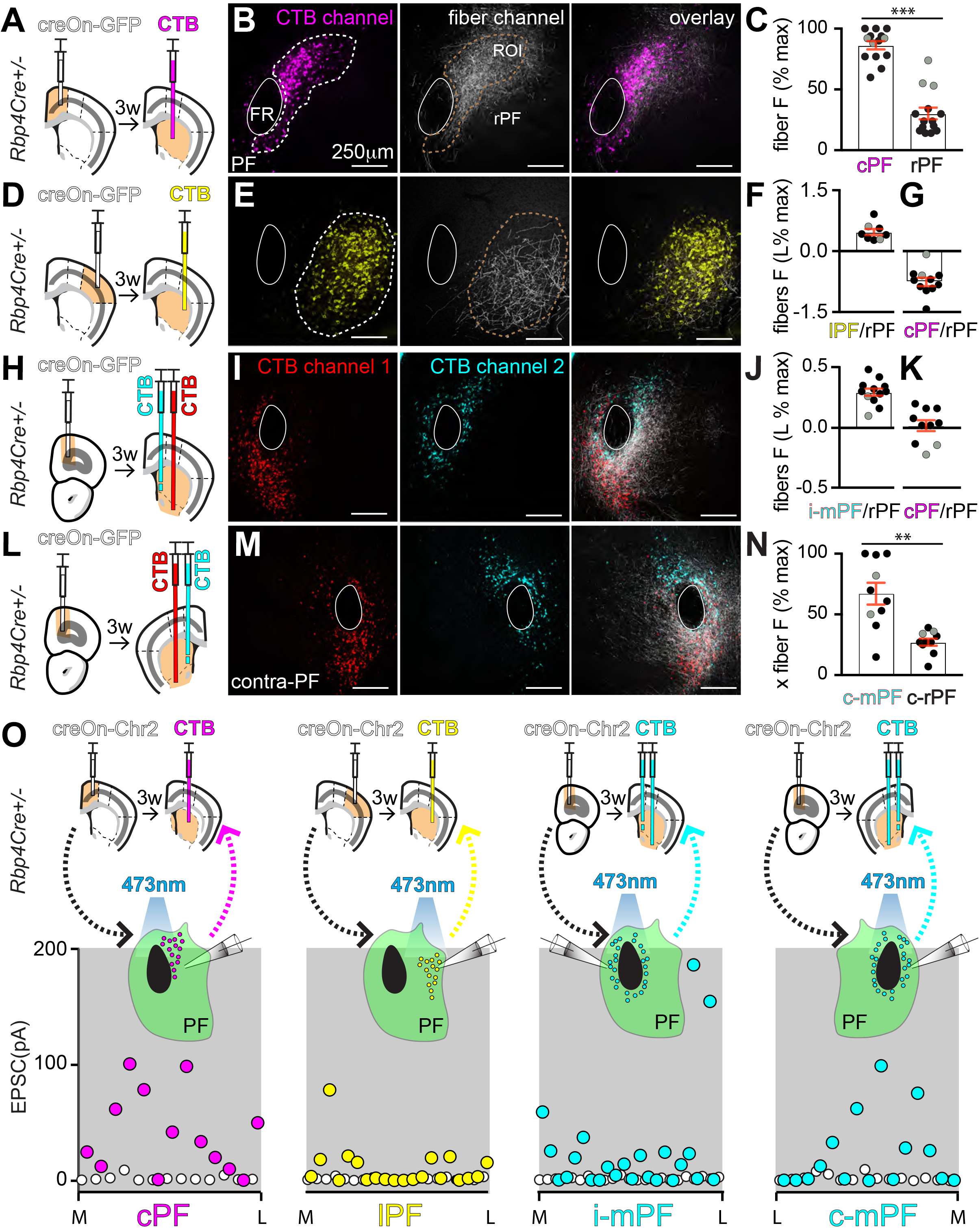
Cortical Layer 5 projections to PF are topographically organized and form closed Cortex-PF-Striatum circuits. **A**, Schematic of coronal sections at +0.9 mm from a *Rbp4-Cre^+/-^* mouse depicting injection of creOn-gfp (white) into layer 5 of secondary motor CTX (MOs) followed by a CTB injection (magenta) into dmSTR 3 weeks later. **B**, Coronal section at -2.1mm in PF showing the results of the experiment in Panel (A). CTB (*left*, magenta) and GFP-expressing axons from MOs (*center,* white) are seen to overlapping in cPF (*right*, overlay). The ROI in the CTB channel (white dashed line) was manually drawn and applied to the GFP channel (brown dashed line) to measure the fluorescence distribution. **C**, Quantification of percentage of the maximal fluorescence intensity (FI) of GFP labeled axons in the cPF ROI, as shown in Panel (B), compared to that in the rest of PF (rPF). Grey filled circles here (and throughout the figure) represent the analyses of coronal section -2.3mm in PF. (n=12/3; slices/animals). P=0.0001; Wilcoxon test. **D-E**, As in (A-B) but with injection of creOn-GFP into primary sensory CTX (SSp) (white) and an injection of CTB (yellow) into dlSTR. **F**, *left,* Quantification of percentage of the maximal FI of GFP labeled axons in the lPF ROI, as shown in panel (E), compared to that in the rest of PF (rPF) in log scale. (n=8/2; slices/animals). **G**, Quantification as in panel (F) but for an experiment with an injection of creOn-GFP into primary SSp and an injection of CTB (yellow) into dmSTR and not dlSTR enabling expression of CTB in the cPF➔dmSTR projections. (Experimental design not shown). This confirmed the specificity of SSp➔lPF fiber topography (n = 11/2; slices/animals). **H-J**, As in panel (D-F) but for injection of creOn-GFP into PFC (white) and CTB injection into mSTR (cyan) and ACB (red). (n = 12/2; slices/animals). **K**, As panel (G) but with an injection of creOn-GFP into PFC and CTB into dmSTR. This confirmed the specificity of PFC➔mPF fiber topography (n = 10/2; slices/animals). **L-N**, As panel (A-C) but for injection of creOn-GFP into PFC and CTB into mSTR and ACB contralateral to injection in PFC. P=0.002; Wilcoxon test. (n = 10/2; slices/animals). **O**, *top,* Schematics of four experimental paradigms using *Rbp4-Cre+/-* mice indicating the sites of injection of creOn-ChR2 into CTX and of CTB into STR three weeks later. Acute brain slices were cut (bottom) and ChR2-stimulated corticothalamic EPSCs were measure at -70 mV in CTB+ neurons in the cPF (n=13/2; cells/mice), lPF (n=13/2), mPF ipsilateral (i-mPF n=23/3) and contralateral (c-mPF n=15/2) to the cortical injection. The amplitude of the ESPC (open circles) and equivalent analysis during a baseline period (closed circles) are shown in previous figures. Grey filled circles in C, F, G, J, K, N represent the analyses of coronal section -2.3mm in PF. Data represented as mean (bar) and s.e.m (red). Also see related figures S7, and S8.

Similarly, analysis of Primary Somatosensory Cortex (SSp) and lPF➔dlSTR PF projection neurons (Figure 7D) (max FI of CTB in: lPF=60%±13; rPF=6%±1; N=8/2; slices /mice; Figure 7E, S7B) revealed overlap of SSp axons and CTB labeled lPF neurons (log ratio of fiber FI in the lPF/rPF = 0.47±0.07; n=8/2; slices/mice; Figure 7E-F; S7B). Conversely, selective targeting of SSp and cPF revealed no overlap between the CTB+ cells and fibers confirming the specificity of SSp➔lPF fiber topography (log ratio of fiber FI in the cPF/rPF=-0.76±0.10; N=11/2; slices /mice; Figure 7G).

Targeting PFC and mPF➔mSTR (Figure 7H) (max FI of CTB in: mPF=75%±7; rPF=18%±1; n=12/2; slices /mice; Figure 7I; S7C) revealed overlap of PFC axons and CTB labeled mPF neurons. (log ratio of fiber FI mPF/rPF=0.29±0.03; N=12/2; slices /mice; Figure 7I-J; S7C). In contrast, PFC axons had little overlap with cPF➔dmSTR PF projection neurons confirming the specific overlap between PFC fibers➔mPF (log ratio of fiber FI in the cPF/ rPF=0.02±0.04; n=10/2; slices /mice; Figure 7K).

Lastly, fibers from PFC also overlapped with mPF➔mSTR PF projection neurons contralateral (con) to the injection site in CTX (Figure 7L) (max FI of CTB: con-mPF=64%±9; rPF=12%±2; n=10/2; slices/mice; Figure 7M; S7D; For max fiber FI overlap in: con-mPF=67%±9; con-rPF= 27%±2; n=10/2; slices /mice; Figure 7M-N; Figure S7D) compared to MOs and SSp which sparsely projected to con-PF in comparison to PF ipsilateral (ipsi) to the injection site (max fiber FI: MOs➔ipsi-PF=89%±2% vs MOs➔con-PF=7%±0.5%; n=2 mice; and SSp➔ipsi-PF=77%±6% vs SSp➔con-PF=2%±0.3; n=2 mice; and PFC➔ipsi-PF=75%±6% vs PFC➔con-PF=30%±3%; n=2 mice; S8).

Pyramidal tract (PT) cells originating in layer 5 project widely throughout the brain and form excitatory synapses onto many classes of neurons but also course through regions without forming synapses (Harris and Shepherd, 2015) (Levesque et al., 1996) Shepherd, 2013). To examine the second question and determine whether the topographically organized PT axons observed in PF (Figure 7A-N; S7, S8) form functional synapse onto PF➔STR projecting cells, whole-cell voltage-clamp recordings were obtained from CTB+cells in PF and axons of layer 5 Cre+ PT cells expressing ChR2 were stimulated. Excitatory synaptic currents were observed in PF➔STR neurons for all the CTX➔PF projections tested (Projections: MOs➔cPF=11/13 EPSCs; n=2;mice; SSp➔lPF=7/21; n=2;mice; PFC➔ipsi-mPF=12/23; n=3;mice; PFC➔con-mPF=7/15; n=2;mice; Figure 7O). Thus, PF striatal projection neurons are a hub for limbic, associative, and sensory-motor information transfer from cortex to STR. Furthermore, each PF subregion targets the same cortical areas from which it receives input, creating circuits organized into cortical-PF-cortical and cortical-PF-striatal motifs. (See discussion for more details).

### Differential modulation of STR by PF sub-classes

Previous work compared inputs from PF➔STR with other thalamic inputs to STR (Ellender et al., 2013) (Alloway et al., 2014). We find that PF has multiple classes of cells that target functionally distinct regions of STR and CTX, and receive disparate inputs from CTX in addition to inputs from the midbrain. There is no connectivity between classes of PF projection neurons, therefore PF forms parallel streams of input to STR, integrating midbrain information with cortical input. However, each channel may have distinct effects on STR by differentially targeting direct and indirect pathway striatal projection neurons (SPNs) and striatal interneurons. To determine if the PF subclasses differently innervate interneurons in STR, whole-cell recordings were made from either cPF➔dmSTR or lPF➔dlSTR projections in Tg(Lhx6-EGFP)BP221Gsat BAC transgenic mice (link: <u>GENSAT resource</u>) (in short, *Lhx6-EGFP*) which expresses GFP in low threshold spiking (LTS) and fast spiking (FS) interneurons and leaves SPNs unmarked (Gittis et al., 2010) (Figure 8A,C). PF to SPNs connectivity was similarly high for both cPF➔dmSTR and lPF➔dlSTR projections (cPF➔dmSTR=9/15 SPNs; n=3 mice; lPF➔dlSTR=9/17; n=4 mice; Figure 8B,D). However, in the same animals, PF connectivity to FS cells, identified based on their firing patterns and membrane properties (Saunders et al., 2016), was low between cPF➔dmSTR (2 of 23 FS cells innervated) compared to lPF➔dlSTR (9 of 14 FS cells innervated) (Figure 8B,D). Inputs to LTS cells, also identified based on their firing patterns and membrane properties was low from both cPF (0 of 11 LTS cells innervated) and lPF (3 of 26 LTS cells innervated) (Figure B,D). Interneuron connectivity between mPF➔mSTR was not examined; however, mPF to SPN connectivity was similarly high (24 of 33 SPNs; n=4 mice; Figure 4H) compared to cPF➔dmSTR and lPF➔dlSTR SPN connectivity. Thus, we find that topographically defined PF➔STR projections all robustly target SPNs but differently innervate LTS and FS interneurons of the STR.

**Figure 8:**
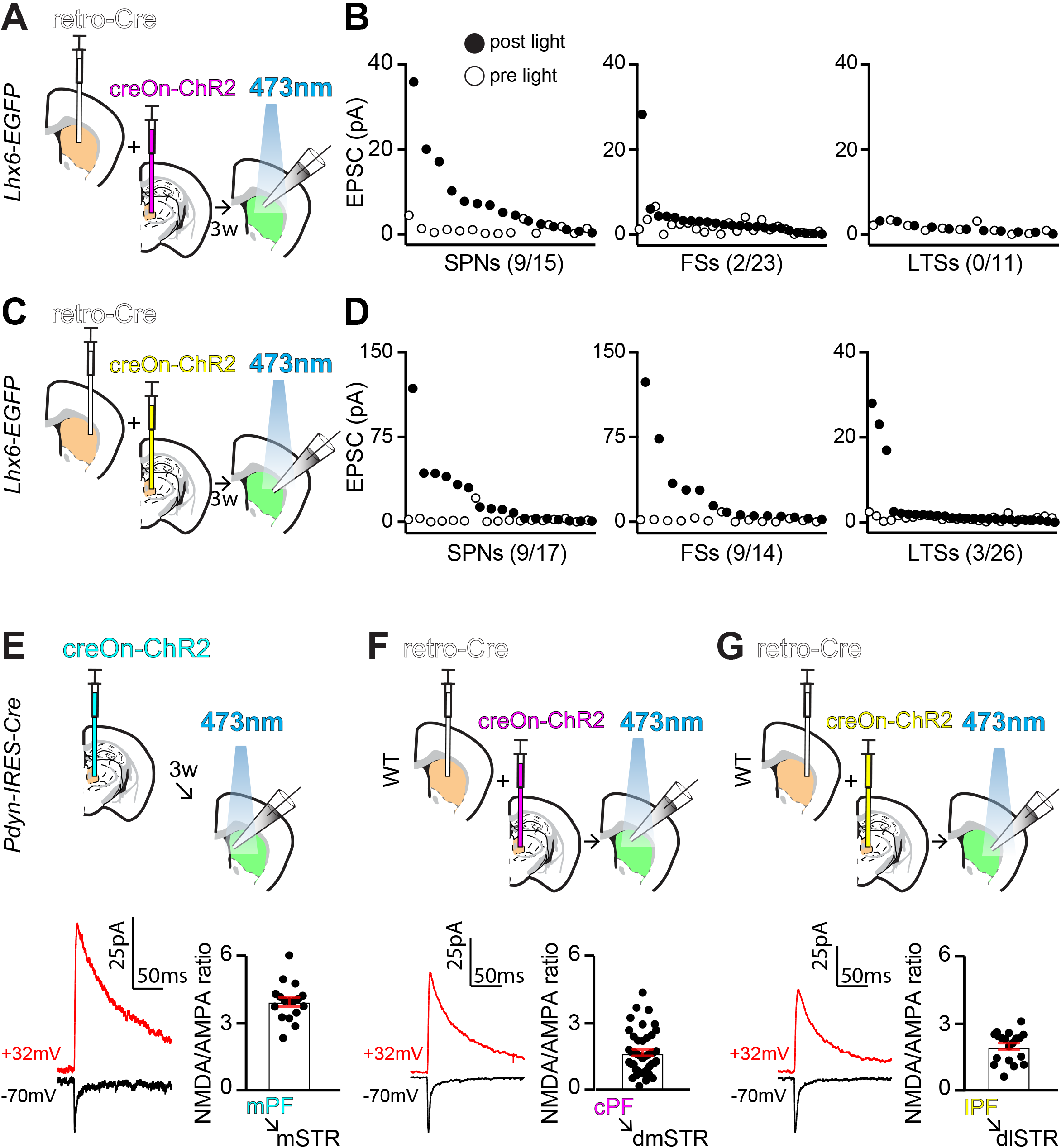
Differential modulation of STR by PF sub-classes. **A**, Experimental design shown of coronal sections at +0.9 mm from a *Lhx6-EGFP* mouse injected with retro-Cre (white) in dmSTR and creOn-chr2 (magenta) in PF. 3 weeks later acute brain slices were cut and ChR2-stimulated cPF➔dmSTR EPSCs were measure at -70 mV. **B**, EPSC amplitudes from SPNs (*left*), FSs cells (*middle*), and LTSs (*right*) in dmSTR. The amplitude of the ESPC (open circles) and equivalent analysis during a baseline period (closed circles) are shown. (n = 3 mice). **C-D**, As in panel (A-B) but with injections of retro-Cre (white) in dlSTR and creOn-ChR (yellow) in PF. (n = 4 mice). **E**, Top: Experimental design shown of coronal sections at -2.1mm from a *Pdyn-IRES-cre* mouse injected with creOn-Chr2 in PF. 3 weeks later whole cell recordings were obtained in STR (highlighted in green). EPSCs were recorded at -70mV and at a holding membrane potential of +20mV from the reversal potential of each cell following a light pulse. Bottom left: representative traces of NMDA (red) and AMPA (black) EPSCs. Bottom right: summary data. (n = 17/2; cells/mice). **F-G**, Same as in panel (E) but for injection of retro-Cre in dmSTR (F) or dlSTR (G) followed by injection of Chr2 into PF.

The NMDA-receptor (NMDAR) component of the SPN glutamatergic EPSC is thought to mediate induction of plateau potentials (up-states) in SPNs (Plotkin et al., 2011). Previous studies report widely varying NMDAR to AMPA-type (AMPAR) glutamate receptor current ratios at PF to SPN synapses: analyses in mice describe that the CTX➔STR synapses induce relatively high NMDAR/AMPAR current ratios compared to the TH➔STR synapses (Ding et al., 2008) whereas the opposite result has been described in rats (Smeal et al., 2007). We reasoned that these differences might have reflected different in the subregions of PF➔STR projections studied as opposed to true interspecies differences. Therefore, we measured AMPAR- and NMDAR-mediated synaptic currents (see methods) for the 3 PF➔STR projections (Figure 8E). The NMDAR/AMPAR currents ratio were higher at mPF➔mSTR synapses (3.8±0.2; n=17/2; cells/mice; Figure 8E) compared to cPF➔dmSTR (1.6±0.1; n=43/4; cells/mice; Figure 8F) and lPF➔dlSTR (1.9±0.2; n=19/3; cells/mice; Figure 8G) which did not differ from one another. Thus, the characteristics of PF➔STR excitatory synapses onto SPNs depend on the target region within the STR.

## DISCUSSION

Here we present a comprehensive cellular and circuit analysis of the PF, a major subcortical excitatory input to the STR. Based on PF’s anatomical, transcriptional, electrophysiological, and synaptic properties we place its projection-neurons into 3 classes of cells. mPF neurons expressed *Pdyn*, the precursor protein for the K-opioid receptor agonist, project to matrix compartments of mSTR and to limbic CTX (for example, ILA, Aid, and PL) and receive bilateral input from layer 5 in PFC. mPF➔STR projection neurons have higher input resistance, lower capacitance, and higher resting potential relative to those in central and lateral aspects of the PF. In cPF, neurons express *Tnc*, project to dmSTR and to limbic and associative regions of CTX (for example, to ILA, MOs, and GU), and receive input from layer 5 associative areas (MOs). Lastly, in lPF, neurons express *Spon1,* project to dlSTR and predominantly sensorimotor regions of CTX (for example, SSp and SSs), and receive input from layer 5 of sensorimotor CTX (SSp). All cell classes in PF have high connectivity to striatal projection neurons but differ in their innervation of striatal interneurons. PF neurons do not interconnect across regions, suggesting that PF subregions do not intermix their incoming cortical and midbrain signals through local inhibitory or excitatory connectivity. Indeed, cells in PF also receive inputs from the SNr, an output structure of the BG, in addition to receiving input from the Superior Colliculus.

### Comparing the mouse PF to that of other species

The anatomical organization that we describe for mouse PF➔STR appears present in other species (Giménez-Amaya et al., 2000) (Jones, 2007). PF/CM in primates may also be separated into 3 regions that preferentially innervate motor CTX (lateral CM), sensory motor STR (medial CM), and associative limbic STR (PF) (Sadikot and Rymar, 2009). In primates CM and PF can be distinguished based on cell density and size (Jones, 2007), vulnerability to disease (Henderson et al., 2000b), as well as *in vivo* firing patterns (Matsumoto et al., 2001) and have been proposed to have different functions (Glimcher and Lau, 2005) (Smith et al., 2014). In rats, lPF projects to dlSTR and mPF projects to dmSTR (Berendse and Groenewegen, 1990) which is a similar, albeit a simplified version of the relationship observed between CM, PF and STR in primates. Nevertheless, no region in the rat TH has been defined as CM due to lack of the lack of a clear histological boundary.

To our knowledge no previous characterization of PF has been conducted in the mouse. It is widely accepted that the gross nuclear division of mouse TH is similar to that in rats (Jones, 2007), although some thalamic nuclear boundaries are more obscure in mice, likely due to a diffused cytoarchitecture. Here, we define 3 classes of neurons in PF as well as functional, transcriptional, and anatomical differences across the medial-lateral aspect of the PF.

The facility of analysis in mice allowed us to uncover differences between subdivisions of the PF that have not been addressable in traditionally genetically-intractable species such as primates, cats, and rats. Even within mouse studies (Parker et al., 2016) (Kato et al., 2011) (Aceves Buendia et al., 2017) (Assous et al., 2017) (Choi et al., 2018) PF has been treated as cellular homogenous and not having subcircuits nor it being distinct to its neighboring TH nuclei. Our findings reveal transcriptional distinctions that demarcate PF cell-type classes and also separate it from its neighboring medial dorsal nucleus in the anterior-posterior axis and the posterior nucleus in medial-lateral axis. Thus, these results permit targeted analyses of specific PF subcircuits and neuron classes in normal behavior and disease-models, similar to studies already underway in other brain regions (Svoboda and Li, 2018) (Beyeler et al., 2018) (Wallace et al., 2017) (Saunders et al., 2015) (Girasole et al., 2018) (Mastro et al., 2014) but not previously possible in the ILM of the TH.

### PF-cortical interactions

Thalamic nuclei typically form reciprocal connections with CTX by receiving input from and projecting to a single cortical region or receiving input from one region and project to another (Sherman, 2016). Thalamic nuclei receive modulatory-inputs from layer 6 while higher-order thalamic nuclei also receive inputs from layer 5 (Harris and Shepherd, 2015) (Sherman, 2016). These CTX-TH-CTX circuits have been proposed to have two functions. First, via recurrent excitation they maintain persistent activity in CTX, as has been shown for projections from motor TH to the anterior lateral motor region of CTX (Guo et al., 2017) and is thought to be necessary for working memory (Bolkan et al., 2017) (Halassa and Kastner, 2017). Second, other CTX-TH-CTX circuits have a triangular motif in which a cortical region targets a second cortical area and a thalamic nucleus that also projects to the second cortical region. This motif, for example, is seen in Pulvinar nucleus outputs to visual cortex, and has been proposed to transmit and synchronize signals about attentional priorities between directly connected cortical regions (Saalmann et al., 2012).

These canonical principals of organization have not been examined fully in ILM TH and its interactions with CTX. It is likely that all subclasses of PF neurons are innervated by layer 6 cortical neurons (Jeong et al., 2016). We find that PF receives input from neurons in layer 5 of limbic, associative, or sensory-motor regions while also projecting back to those regions. Thus, it seems that the laminar sources of cortical input to PF are similar to those for other nuclei of TH.

PF➔CTX projections align with CTX➔PF projections, suggesting the existence of a corticothalamic recurrent network through PF. However, unlike typical TH nuclei, the main output of PF is to STR and not to CTX. For this reason, and analogous to the triangle attentional motif described above, we propose that cortical-PF-cortical circuits facilitate and shape striatal output of a behaviorally relevant cortical region. In parallel, PF can integrate information from CTX with that from subcortical nuclei (such as SC and SNr) to facilitate correct action selection in an ongoing sensorimotor context. Since we find that PF neuron classes are not interconnected in PF, these networks of activity can act relatively independently of each other.

We find that PFC projects bilaterally to mPF whereas SSP➔lPF or MOs➔cPF projections are strictly ipsilateral, highlighting the potential different functions of the PF subclasses characterized here – laterality may be important to maintain for sensory and motor circuits while perhaps a global limbic signal may need to be dispersed across both hemispheres. PT cortical neurons do not project to contralateral STR (Harris and Shepherd, 2015) thus the bilateral PFC➔mPF projections may transmit PT related-activity signal near synchronously to both STR without recruitment of bilateral IT-type cortical projections.

## METHODS

### Mice

This study is based on data from mice postnatal day 50, both males and females. We used the C57BL/6NCrl (Charles River Laboratories, Wilmington, MA, stock #027) and transgenic mice lines: *Dyn-IRES-cre* (Jackson Laboratories, Bar Harbor, ME, stock #027958), Rbp4-Cre (Gensat project, founder line KL100), and *LHX6-EGFP* (Gensat project, founder line BP221). Animals were maintained on a C57BL/6 background and kept on a 12:12 light/dark cycle or a reversed cycle under standard housing conditions.

Experimental manipulations were performed in accordance with protocols approved by the Harvard Standing Committee on Animal Care following guidelines described in the US National Institutes of Health Guide for the Care and Use of Laboratory Animals. All mice brain coordinates in this study are given with respect to Bregma; anterior–posterior (A/P), medial–lateral (M/L), and dorsal–ventral (D/V).

### AAVs

Recombinant adeno-associated viruses (AAVs of serotype 1,2,8 or 9) encoding a double floxed inverted (DFI) gene under the control of CAG, Ef1a, or hSyn promotors were used to express the gene in the Cre-recombinase expression neurons. Additionally, we used intersectional AAVs that expressed the gene only when Flp is present and Cre is absent (FlpOn/CreOff). Retrograde AAVs which efficiently infect axons (Tervo et al., 2016) were used to deliver Flp or Cre recombinase to neurons upstream to the injection site. AAVs were packaged by commercial vector core facilities (UNC Vector Core and Penn Vector Core) and upon arrival stored at a working concentration 10^11 to 10^13 genomic copies per ml at -80C.

### Rabies viruses

Rabies viruses carrying the transgene for the H2B:EGFP fusion protein were generated in-house. Synthesized H2B-EGFP vector was cloned into pSPBN-SADΔG-tdTomato plasmid using SmaI and NheI restriction sites, replacing a tdTomato sequence. B19G-SADΔG-H2B:EGFP virions were first generated via cDNA rescue using a procedure based on previously described protocols (Wickersham et al., 2010). Briefly, HEK 293T cells (ATCC CRL-11268) were transfected with pSPBN-SADΔG-H2B:EGFP, pTIT-B19N, pTIT-B19P, pTIT-B19G, pTIT-B19L and pCAGGS-T7 using the Lipofectamine 2000 transfection reagent. 5 to 7 days post-transfection, the supernatant was filtered through a 0.22 μm PES filter and transferred to BHK-B19G cells for amplification. Virions were then serially amplified in three rounds of low-MOI passaging through BHK-B19G cells by transfer of filtered supernatant, with 3 to 4 days between passages. Cells were grown at 35 °C and 5% CO_2_ in DMEM with GlutaMAX (Thermo Scientific, #10569010) supplemented with 5% heat-inactivated FBS (Thermo Scientific #10082147) and antibiotic-antimycotic (Thermo Scientific #15240-062). For concentrating the virions, media from dishes containing virion-generating cells was first collected and incubated with benzonase nuclease (1:1000, Millipore #70664) at 37°C for 30 min before filtering through a 0.22 μm PES filter. The filtered supernatant was transferred to ultracentrifuge tubes (Beckman Coulter #344058) with 2 ml of a 20% sucrose in dPBS cushion and ultracentrifugated at 20,000 RPM (Beckman Coulter SW 32 Ti rotor) at 4°C for 2 hours. The supernatant was discarded and the pellet was re-suspended in dPBS for 6 hours on an orbital shaker at 4 °C before aliquots were prepared and frozen for long-term storage at -80 °C. Unpseudotyped rabies virus titers were estimated based on a serial dilution method (Osakada & Callaway, 2013) counting infected (H2B:EGFP^+^) HEK 293T cells, and quantified as infectious units per ml (IU/ml).

B19G-SADΔG-EGFP, EnvA-SADΔG-EGFP, and B19G-SADΔG-ChR2-EYFP viruses were generated by amplification from existing in-house stocks using similar passaging procedures described above. Pseudotyping was performed after the last passaging round of unpseudotyped virion amplification. BHK-EnvA cells were infected with the filtered supernatant containing unpseudotyped virions for 6 hours, followed by two rounds of trypsinization with dPBS washes and re-plating over two consecutive days. Pseudotyped rabies virus titers were estimated as described above counting infected (EGFP^+^) HEK 293T-TVA800 cells. For quality control, pseudotyped rabies virus stocks were tested *in vitro* for leak of unpseudotyped virus with a similar titering protocol by infecting HEK 293T cells. Virus batches used had a leak of less than 2 × 10^3^ IU/ml.

### Stereotaxic Intracranial Injection

Mice were anesthetized with 2.5% isoflurane in 80% oxygen and placed in a stereotaxic frame (David Kopf Instruments Model 900). Under aseptic conditions, the skull was exposed and leveled (<100 μm difference between 1.5+A/P and Lambda as a cutoff for proper leveling of the skull). 250 μm craniotomies were made with an electric drill (Foredom Electric Company K.1070) with a ball bur (Busch and Co. S33289) attached to the manipulator. All reagents were injected through a pulled glass pipette (Drummond Scientific Company pipettes) with a tip of approximate 50 μm (pulled with a P-97 model Sutter Instrument Co. pipette puller). To avoid leak into other brain regions and back spill through the pipette track the injection pipette was lowered 200 μm ventral to the region of injection before being brought up to the point of injection. The pipette was left in place for 3min prior to injection and the reagent of interest was delivered at a rate of 50nl/min using a UMP3 micro-syringe pump (World Precision Instruments). Following injection, we waited 5min at the injection site before raising the pipette 200 μm above the injection site and waited 5 minutes more. We then exited the brain at about 1mm/minute. To minimize their dehydration during surgery mice received a subcutaneous injection of 1ml of sterile saline (Teknova S5819). Additionally, in order to reduce inflammation, mice received an injection of Ketoprofen (Zoetis 07-803-7389) at an amount of 0.01mg per gram of animal mass. Postoperatively, mice were monitored on a heat pad for one hour before being returned to their home cage. Mice were then monitored daily for at least 5 days and received a MediGel Carprofen cup in their home cage (Clear H_2_O).

### Injection coordinates

All coordinates that were used in this study were relative to Bregma. For PFC; 2.8 mm A/P, 1.2 mm M/L, 0.9 mm D/V, MOs; 0.8, 0.9, both 0.8 and 0.5, SSp; 1.0, 2.2, 1.0, mSTR; 0.8, 1.0, both 3.3 and 2.7, dmSTR; 0.8, 1.6, 2.6, dlSTR; 0.8, 2.4, 2.5, ACB; 0.8, 1.5, 4.6, mPF; -2.1, 0.5, both 3.7 and 3.5, lPF; -2.1, 0.88, both 3.75 and 3.55.

### Injection volumes and waiting time for specific anatomical regions and reagents

mSTR: RV-nGFP (150-200nl), CTB (80-200); dmSTR: RV-nGFP (150-200), CTB (80-200), retro-Flp (200), retro-Cre (300), RV-GFP (100), RV-ChR2 (200); dLSTR: RV-nGFP (150-200), CTB (80-200), retro-Flp (200), retro-Cre (300), RV-GFP (100), RV-ChR2 (200); mPF: creOn-GFP (75-150), CreOn-ChR2-mCherry (200-300), CreOn-ChR2-GFP (200-300), FlpOn/CreOff (200), CreOn-TVA (100), CreOn-OG (100), p.RV-GFP (150-200); LPF: FlpOn/CreOff (200), CreOn-ChR2-mCherry (200-400); ACB: CTB (80-140); MOs: creOn-GFP (200), CreOn-ChR2-GFP (250); SSp: creOn-GFP (200), creOn-TdTom (200), CreOn-ChR2-mCherry (100-250); PFC: creOn-GFP (200-250), CreOn-TdTom (200-250), CreOn-ChR2-GFP (100-250). Waiting times for reagents were as followed: CTB: 3-7 days; AAVs: 2.5-5 weeks; RV-nGFP: 5-10 day; RV-EGFP: 7-10 days. RV-ChR2: 3-7 days; p.RV-GFP: 5-10 days;

### Histology and Imaging for STPT

Animals were perfused transcardially with ice-cold 0.9 % saline solution followed by 4% paraformaldehyde (PFA) (diluted in 0.2 M phosphate buffer) for 7 min at 7 ml/min, 5 to 10 days after RV-nGFP infection. Brain were fixed in 4% PFA for 24h before being transferred to 0.1 M glycine solution (diluted 0.1 M phosphate buffer), for 48h at 4 °C before being stored in 0.1 M phosphate buffer at 4 °C until imaged. Imaging was done as previously described (Ragan et al., 2012). In short, brains were embedded in 4% agarose in 0.05M PB, cross-linked in 0.2% sodium borohydrate solution (in 0.05 M sodium borate buffer, pH 9.0-9.5). The entire brain (including the olfactory bulb and the cerebellum) was imaged with a high-speed 2-photon microscope with integrated vibratome at 1μm-1μm x-y resolution for a depth of 50 μm on a TissueCyte 1000 (TissueVision). The 2-photon excitation wavelength was 910 nm, which efficiently excites GFP. A 560 nm dichroic mirror (Chroma, T560LPXR) and band pass filters (Semrock FF01-520/35 an) were used to separate green.

### Histology and imaging for all other experiments

mice were anesthetized with isoflurane and perfused transcardially with 4% PFA in 0.1 M sodium phosphate buffer (PBS). Brains were post-fixed for 24-48 hours and transferred to a 0.1M PBS solution until further processing. Coronal slices were made at 50 μm thickness per slices on a vibrating blade microtome (Leica Biosystems VT1000S). Brain sections were mounted on superfrost slides (VWR 48311-703) dried, and cover-slipped with ProLong antifade reagent containing DAPI (ThermoFisher P36962). Whole slides were imaged with an Olympus VS120 slide scanning microscope with a 10X objective. Regions of interest were imaged with an Olympus FV1200 confocal microscope using a 10X or 60X objectives at the Harvard Neurobiology Imaging Facility and Harvard Neurodiscovery Imaging Core.

### Immunohistochemistry

Images requiring immunohistochemical staining were processed using a protocol previously described in Pisanello and Mandelbaum et al. (Pisanello et al., 2017). In short, slices were incubated in PBS blocking solution containing 0.3% Triton X-100 (PBST) for 1h at RT (20–22 °C). Slices were then incubated over night at 4 °C in the same blocking solution with 1% goat serum with MOR primary antibody (Life Technologies AB5511). The next day, slices were rinsed 3 × 10 min in PBS before being incubated in the blocking solution with secondary antibody (1 mg/mL goat anti-rabbit Alexa Fluor 647 (AB_2535812; Life Technologies) or Alexa Fluor 594 (R37117; Life Technologies) Life Technologies). The slices were then rinsed again, mounted, and imaged as described above.

### In situ hybridization

Tissue for in situ hybridization was processed using a protocol previously described in Hvartin and Hochbaum et al. (Hrvatin et al., 2018). In short, animals were euthanized and brains were immediately frozen on dry ice to be sliced via cryostat (Leica CM 1950). Cells in the mPF were marked by *ProDynorphin*, cells in cPF by *TNC*, and cells in lPF by *Spon-1*. Excitatory cells in TH were marked with *Slc17a6* (Vglut-2). For the image presentation of the ISH in Figure 3, nuclei masks were created and each nucleus was pseudo-colored according to the number of puncta contained within the specific mask, as previously described in Hvartin and Hochbaum et al. (Hrvatin et al., 2018).

### STPT Image Analysis

Raw images were corrected for non-uniform illumination and light collection, stitched in 2D, and stacked in 3D. **RV-nGFP cells counts.** GFP+ neurons were automatically detected by a convolutional network (ID: 164) trained to recognize nuclear neuronal cell body labeling. The 3D stack was then registered to a 3D reference brain based on the ABA (Kim et al., 2015) (Sunkin et al., 2013) by 3D affine registration followed by a 3D B-spline registration using the software Elastix (S.Klein et al., 2010). The number of total input neurons in each brain region was normalized by the total number of GFP+ cells detected in the parent region. (For example, in Figure 1D the parent region is sub-CTX).

### Brain volume quantification

To measure the volume of anatomical regions, the average reference brain (built using 40 STPT imaged brains) was aligned to the ABA. Segmentation areas were registered onto each brain using the b-spline registration procedure described above (i.e. the ABA segmentation was registered onto each individual brain). The number of voxels that belong to each region in the transformed ABA segmentation were counted and multiplied by 0.02 × 0.02 × 0.05 mm^3^ (the dimensions of an anatomical voxel unit), resulting in the total volume of each region. **Projection mapping data processing.** Previously published methods were adopted for quantifying neuronal projections as imaged by STPT (Oh et al., 2014). In brief, filtered images of the original image data were generated by applying a square root transformation, histogram matching to the original image, and median and Gaussian filtering using ImageJ (NIH) software. The original images were then subtracted from the filtered images to generate signal images. These were then converted to binary maps by applying a threshold chosen to maximize signal retention while minimizing background auto-fluorescence. We cannot rule out the possibility that faint and sparse signals were being missed in our automatic detection. False-positive signals at the injection sites and from bright fluorescence from the dura were removed using manually curated masks for each brain. The method measures fluorescence from all axons, including axons of passage. For this reason, we only analyzed signals in CTX, where fibers of passage are less likely. To calculate the putative output of PF to CTX the relative axon density was measured as fraction of all GFP+ pixels that are located in a given area divided the fraction of cortical volume contained in the area. This metric gives the relative enrichment of axons in each cortical area compared to a uniform distribution of axons within CTX. The scalable brain atlas (<u>link</u>) was used for Figure 6 C-H visualization (Bezgin et al., 2009).

### Image analysis for all other experiments

Quantification of the fluorescence intensity (FI) of CTB+ cells in PF were done using a custom macro in ImageJ (NIH). For each coronal section in PF, and based on the ABA, the mean pixel FI was calculated across a ventral-dorsal line with an 0.6 μm medial-lateral width. Using Graph-Pad prism (GraphPad Software, La Jolla, CA), a 2^nd^ order smoothing (Savitzky and Golay, 1964) was applied with 200 nearest neighbors prior to normalizing each channel. For quantification of the GFP+ cell bodies in PF, and GFP+ axons in STR (Figure 4) the same analysis was used, with the exception of the ventral dorsal line being 1.0mm wide. For analysis of *Pdyn*+PF➔STR axons coronal sections of STR at +0.6, +0.9, and +1.2 mm were grouped together. For the analysis of topographical organization of *Pdyn*+ axons in STR (Figure 4) patches (based on MOR stain) were manually labeled while being blinded to the GFP+ PF➔STR fiber location. Using a custom macro in ImageJ each patch label was expanded by 100 μm in all directions and the mean FI of the patch and this peri-patch region were calculated for the MOR channel and the fiber channels. For image analysis of the Layer 5 CTX➔PF inputs (Figure 7) the DAPI channel was used to mark the location of PF to ensure that the selection was done solely based on anatomical location as defined in Figure 1 and based on the ABA. The CTB channel was used to label the region in PF with all CTB+ cells and mean FI was measured in the that region compared to the rest of PF (Figure S7). Next, the areas define as encompassing the CTB+ cells in PF was applied to the CTX➔PF fiber channel and FI of fibers were calculated in that region and compared to the rest of PF. Background FI was calculated by taking the mean of 3 random tissue areas of 0.3mm^2^. The comparison of the FI inputs from CTX to PF, ipsilateral to the injection sight vs. contralateral to the injection sight (Figure S8) was done by manually labeling the axons ipsilateral to the injection sight. The axon location contralateral to the injections was at a similar location and shape as ipsilateral to the injection, allowing to apply the same label to the ipsilateral side. Correction for background FI was done as described above.

### Whole-cell dissociation and RNA capture

Dissociated whole-cell suspensions were prepared using a protocol adapted from Hrvatin amd Hochbaum et al. (Hrvatin et al., 2018). 8-week old C57BL/6NCrl male mice (Charles River Laboratories, Wilmington, MA, stock #027) were pair-housed for a few days after arrival in a regular light/dark cycle room prior to tissue collection. Mice were transcardially perfused with an ice-cold choline cutting solution containing neuronal activity blockers (110 mM choline chloride, 25 mM sodium bicarbonate, 12 mM D-glucose, 11.6 mM sodium L-ascorbate, 10 mM HEPES, 7.5 mM magnesium chloride, 3.1 mM sodium pyruvate, 2.5 mM potassium chloride, 1.25 mM sodium phosphate monobasic, 10 μM (R)-CPP, 1 μM tetrodotoxin, saturated with bubbling 95% oxygen/5% carbon dioxide, pH adjusted to 7.4 using sodium hydroxide). Brains were rapidly dissected out and sliced into 250 μm thick coronal sections on a Leica VT1000 vibratome in a chilled cutting chamber filled with choline cutting solution. Coronal slices containing the TH were then transferred to a chilled dissection dish containing choline cutting solution for microdissection of the PF under a stereomicroscope. Dissected tissue chunks were transferred to cold HBSS-based dissociation media (Thermo Fisher Scientific Cat. # 14170112, supplemented to final content concentrations: 138 mM sodium chloride, 11 mM D-glucose, 10 mM HEPES, 5.33 mM potassium chloride, 4.17 mM sodium bicarbonate, 2.12 mM magnesium chloride, 0.9 mM kynurenic acid, 0.441 mM potassium phosphate monobasic, 0.338 mM sodium phosphate monobasic, 10 μM (R)-CPP, 1 μM tetrodotoxin, saturated with bubbling 95% oxygen/5% carbon dioxide, pH adjusted to 7.35 using sodium hydroxide) supplemented with an additional inhibitor cocktail (10 μM triptolide, 5 μg/ml actinomycin D, 30 μg/ml anisomycin) and kept on ice until dissections were completed. The remaining tissue was fixed in 4% PFA in PBS for histological verification. Dissected tissue chunks from 8 mice were pooled into a single sample for the subsequent dissociation steps. Tissue chunks were first mixed with a digestion cocktail (dissociation media, supplemented to working concentrations: 20 U/ml papain, 1 mg/ml pronase, 0.05 mg/mL DNAse I, 10 μM triptolide, 5 μg/ml actinomycin D, 30 μg/ml anisomycin) and incubated at 34 °C for 90 min with gentle rocking. The digestion was quenched by adding dissociation media supplemented with 0.2% BSA and 10 mg/ml ovomucoid inhibitor (Worthington Cat. # LK003128), and samples were kept chilled for the rest of the dissociation procedure. Digested tissue was collected by brief centrifugation (5 min, 300 *g*), re-suspended in dissociation media supplemented with 0.2% BSA, 1 mg/ml ovomucoid inhibitor, and 0.05 mg/mL DNAse I. Tissue chunks were then mechanically triturated using fine-tip plastic micropipette tips of progressively decreasing size. The triturated cell suspension was filtered in two stages using a 70 μm cell strainer (Miltenyi Biotec Cat # 130-098-462) and 40 μm pipette tip filter (Bel-Art Cat. # H136800040) and washed in two repeated centrifugations (5 min, 300 *g*) and re-suspension steps to remove debris before a final re-suspension in dissociation media containing 0.04% BSA and 15% OptiPrep (Sigma D1556). Cell density was calculated based on hemocytometer counts and adjusted to approximately 100,000 cells/ml. Single-cell encapsulation and RNA capture on the inDrop platform was performed at the Harvard Medical School ICCB Single Cell Core using v3 chemistry hydrogels based on previously described protocols (Zilionis et al., 2017). Suspensions were kept chilled until the cells were flowed into the microfluidic device. The encapsulated droplets were broken and cDNA was processed for next-gen sequencing, as previously described (A. M.Klein et al., 2015) generating index libraries that were then pooled and sequenced across 3 runs on the NextSeq500 (Illumina) platform.

### Acute Brain Slice Preparation and electrophysiology experiments

Experiments were done as previously described (Wallace et al., 2017). In short, artificial cerebrospinal fluid (ACSF) containing 2mM [Ca] and 1 mM [Mg] was superfused at 3 ml/min. For optogenetics experiments, 3 to 5 ms duration light pulses from a 473 nm laser (at 5-10mW per mm^2^ (measured at the sample plane) were used. Recordings were performed at 32C using Cs-based internals for voltage-clamp measurements of synaptic currents and K-based internals for current-clamp measurements of firing patterns.

### In drops analysis

Transcripts were processed according to a previously published pipeline (A. M.Klein et al., 2015) (Hrvatin et al., 2018). Briefly, a custom transcriptome was assembled from the Ensembl GRCm38 genome and GRCm38.84 annotation using Bowtie 1.1.1, after filtering the annotation gtf file (gencode.v17.annotation.gtf filtered for feature_type=”gene”, gene_type="protein_coding" and gene_status="KNOWN"). Read quality control and mapping against this transcriptome was performed using default parameters. Unique molecular identifiers (UMIs) were used to reference sequence reads back to individual captured molecules. The output matrix (cells x genes) was then filtered to exclude cells with less than 500 UMIs and used as the input to the Seurat pipeline for further analysis (Satija et al., 2015). Genes were excluded if UMIs were found in 3 cells or less. Cells were excluded if they expressed fewer than 400 genes, or more than 5500 genes. Cells with 15% or more of their transcriptome derived from mitochondrial genes were excluded. Finally, cell doublets were estimated by creating synthetic doublets from the dataset and computing a k-nearest neighbor graph (k = 30) with both cells and synthetic doublets. Cells were ranked according to the percentage of nearest neighbors that were synthetic doublets. Cells in the top 5% of doublet scores were excluded as putative doublets. Cells were then log-normalized and scaled to 10,000 transcripts per cell. Variable genes were identified using the MeanVarPlot() function, which calculates the average expression and dispersion for each gene, then bins genes and calculates a z-score for dispersion within each bin. The following parameters were used to set the minimum and maximum average expression and the minimum dispersion: x.low.cutoff=0.0125, x.high.cutoff=3, y.cutoff=0.5. Next, the count matrix was regressed against the number of UMIs and percentage of counts comprising mitochondrial genes and scaled. Then PCA was carried out and the top 20 principal components (PCs) were kept. Finally clustering was performed using the FindClusters() routine. Clustering resolution was set to 0.6. This resulted in 13 initial clusters, that were categorized into 7 broad cell-type classes by canonical gene expression patterns (Mrc1/Cd36 for macrophage, Olig1/Pdgfra for oligodendrocytes and oligodendrocyte precursors, Vtn for pericytes, Cldn5/Pecam1 for endothelial and smooth muscle cells, Aqp4 for astrocytes, P2ry12/ Cx3cr1 for microglia, and Snap25/Syn1 for neurons). **Subclustering of neurons.** Cells from neuronal clusters were merged and re-clustered as above with 10 PCA components (estimated as significant by the JackStraw algorithm), yielding 6 initial clusters. Differential gene expression was carried out using Monocle2 (Trapnell et al., 2014). Only 3 clusters had 2-fold enriched genes (Clusters without 2-fold enriched genes were not considered distinct cell types, but instead a result of overclustering). Cells in these 3 clusters were used as a training set to classify the other cells using a random forest classifier (using the Seurat function ClassifyCells()). Bootstrapping by repeating this classification process 1000 times produced a metric for classification. Cells that were classified < 95% of the time to the same cluster were excluded. This final classification was used as input for differential gene expression using Monocle2.

### Electrophysiological Analysis

Electrophysiological properties and ChR2-evoked EPSCs were performed using automated scripts written in MATLAB. Following the electrophysiology analysis white papers of the ABA (link: <u>electrophysiology overview</u> <u>technical whitepaper</u>) we did not make *a priori* assumptions about the input resistance, resting potential, or minimal firing rate necessary to designate a cell as “healthy”. Therefore, our final data set includes neurons that, for example, do not fire any action potentials to injected current. Selection of neurons for inclusion was based on the series resistance. In short, tables containing the name of the cell, annotations about the position of the cell and conditions of the experiment were noted during the experiment including the start and stop sweeps to be analyzed which were then used to automatically retrieve and analyze data. **Intrinsic properties.** Resting membrane potential was measured as the median of potentials during periods of the sweep that had with no current injection. Membrane capacitance (Cm) and resistance (Rm) as well as series resistance were measured in voltage-clamp mode by fitting a single exponential to the current evoked by a -5 or -10 mV voltage pulse. Rs was estimated from the peak of the exponential fit – i.e. Rs= ΔV/ΔI(t=0) with t=0 being the start of the voltage step command. The steady state current was used to calculate Rm=ΔV/ΔI(t=∞) - Rs. Cm is then calculated from the time constant of the fit tau=RsRmCm/(Rs+Rm). **EPSCs.** Resting membrane properties were measured as above. To determine the amplitude of the EPSC and compare it to the amplitudes expected from chance fluctuations of the membrane potential (e.g. due to thermal and seal noise or spontaneous synaptic events), two 15 ms long periods were analyzed in each sweep. The first was a time window after the light pulse in which a genuine ChR2-evoked EPSC would be expected. In this window, the average current compared to baseline was calculated, as well as its peak deviation (positive for NMDA-receptor mediated currents at positive potentials and negative for AMPA-receptor mediated currents at rest). In addition, a “peri-peak” value was calculated from the average current in a time window (3 ms long) around the time of the peak deviation from rest in the average of all sweeps for the cell. Identical analyses were carried out in a time window occurring 50 or 100 ms before the light pulse to estimate baseline fluctuations for each parameter. All measurements of AMPA-receptor mediated EPSCs were done at - 70 mV. For NMDA-receptor mediated EPSCs, the reversal potential of the EPSC was found (typically at nominally +5-10 mV) and the cell was depolarized a further 20 mV, at which the measurement was made. No corrections were made for liquid junction potential (~ 8 mV). FSs, LTSs, and SPNs in the LHX6-GFP were identified based on responses to current injections, membrane resistance, and the presence or absence of dendritic spines as previously described in Saunders et al (Saunders et al., 2016).

### Statistical analyses

Data points are stated and plotted as mean values ± SEM. p values are represented by symbols using the following code: * for 0.01<p<0.05, ** for 0.001<p<0.01, and *** for p<0.001. Exact p-values are stated in figure legends and main text. All statistical tests were non-parametric as noted in the text. No *a priori* power analyses were done.

### Data software and availability

Data and software are available upon request. Table 1 gives a list of all the acronyms used in all the figures. Table 2 gives the data from experimental design shown in Figure 1A. Table 3,4,5,6 gives full genes list from the analysis done in Figure 3. Table 7 gives the full data set for analysis done in Figure 6.

## Supplementary information

**Table 1:** abbreviations lookup table.

**Related to Figure 1:**

**S1:**
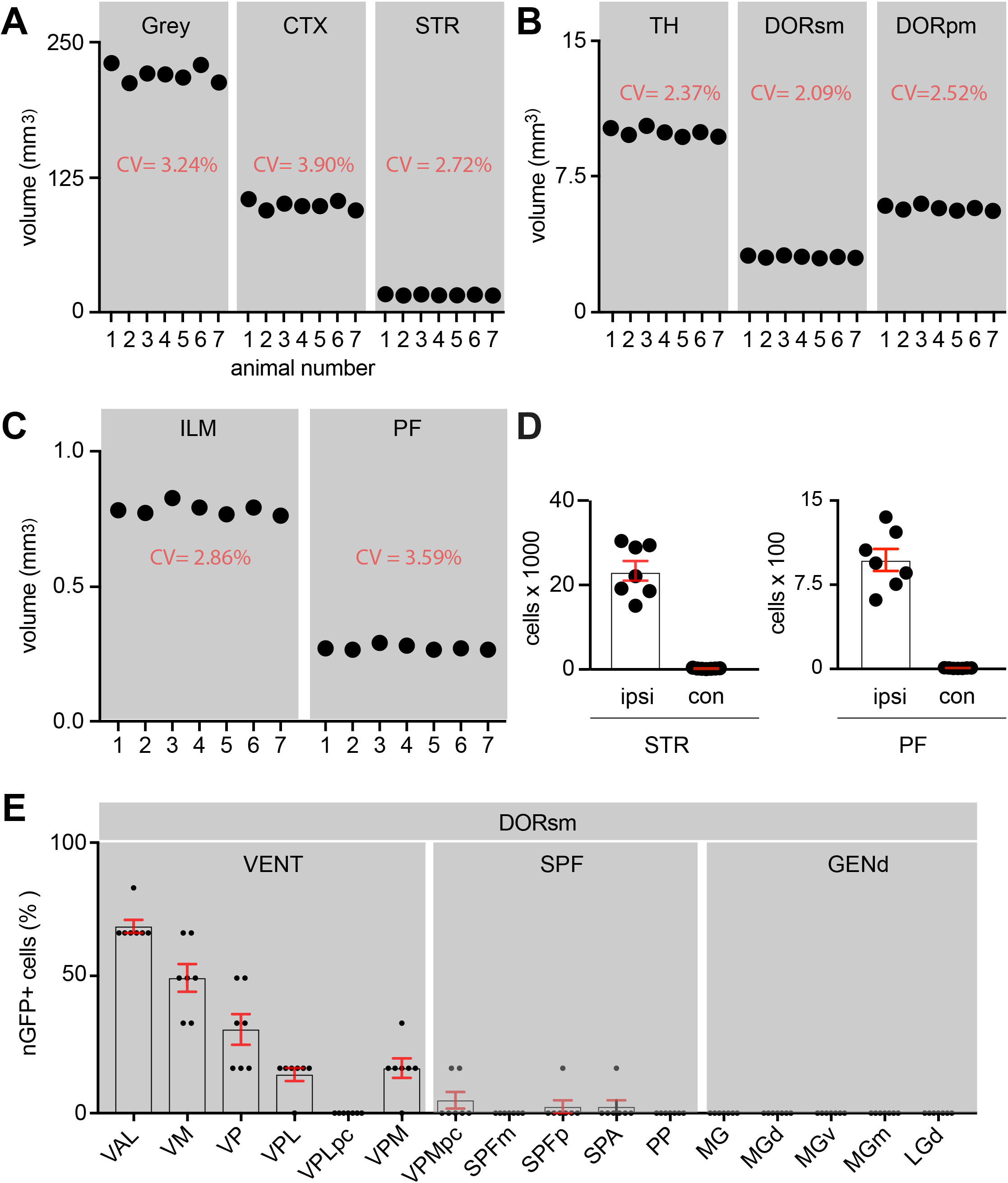
**A-C**, Volume of regions of interest shown for the 7 mice which were pooled together for subsequent analysis. Coefficient of variation (CV) of each region volume across the 7 mice is noted in red. **D**, Number of RV-nGFP+ cells ipsilateral (ipsi) or contralateral (con) to the injection site in STR (*left*) and PF (*right*). (PF: n=6,848/7; STR: n=165,382/7; cells/mice). **E**, Percent of total RV-nGFP+ cells distributed across nuclei-groups and nuclei in sensory-motor cortex related TH (DORsm) (n=32,99/7; cells/mice). **Movie1:** Example of a whole brain from experimental design shown in Figure 1A.

**Table2:** Cell counts and volumes from all brain regions analyzed across 7 mice from the experiment shown in Fig 1A.

**Related to Figure 2:**

**S2:**
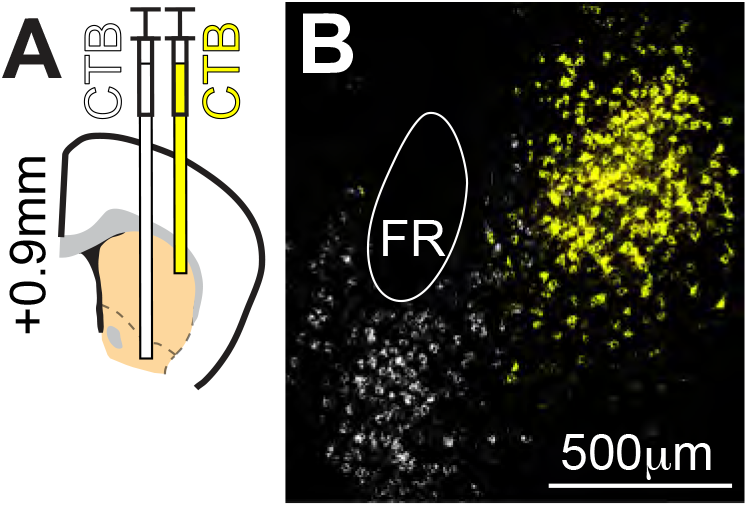
**A**, Experimental design shown by a schematic of a coronal section at +0.9mm from a WT mouse depicting 2 injections of 2 CTB variants in the dlSTR (yellow) and nucleus accumbens (ACB) (white). The injection of the 2 variants of the CTB into two anatomical defined regions allowed to ask whether there is additional topography between PF and STR. The injection site regions (STR and ACB) are highlighted in orange. **B**, An example of a PF coronal section at -2.1mm from the experiment shown in A. The FR is highlighted in the tissue and labeled to help orient to mPF.

**Movie2:** Video of coronal section of tissue clearing in TH starting anterior to PF and going posterior to PF depicting the CTB labeling of the cPF➔dmSTR projection system.

**Related to Figure 3:**

**S3:**
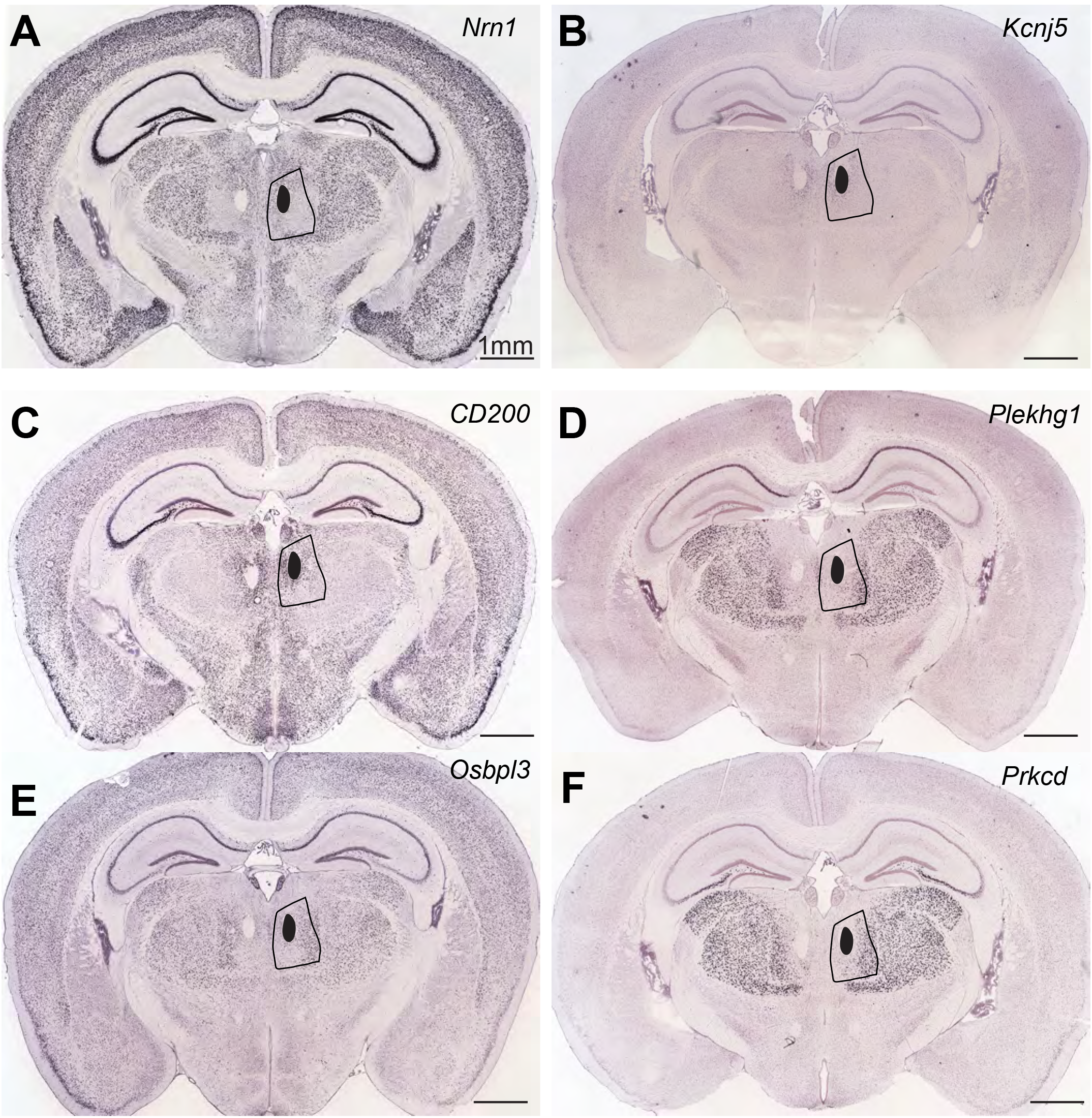
Example of *in situ hybridization* from the ABA with genes that are highly correlated (A-C) or anti-correlated (D-F) with *Pdyn* expression on a cell-by-cell basis.

**Table 3:** list of genes whose expression is elevated in Cluster 1.

**Table 4:** list of genes whose expression is elevated in Cluster 2.

**Table 5:** list of genes whose expression is elevated in Cluster 3.

**Table 6:** list of genes whose expression was found to be correlated or anti-correlated with *Pdyn*.

**Related to Figure 4:**

**S4:**
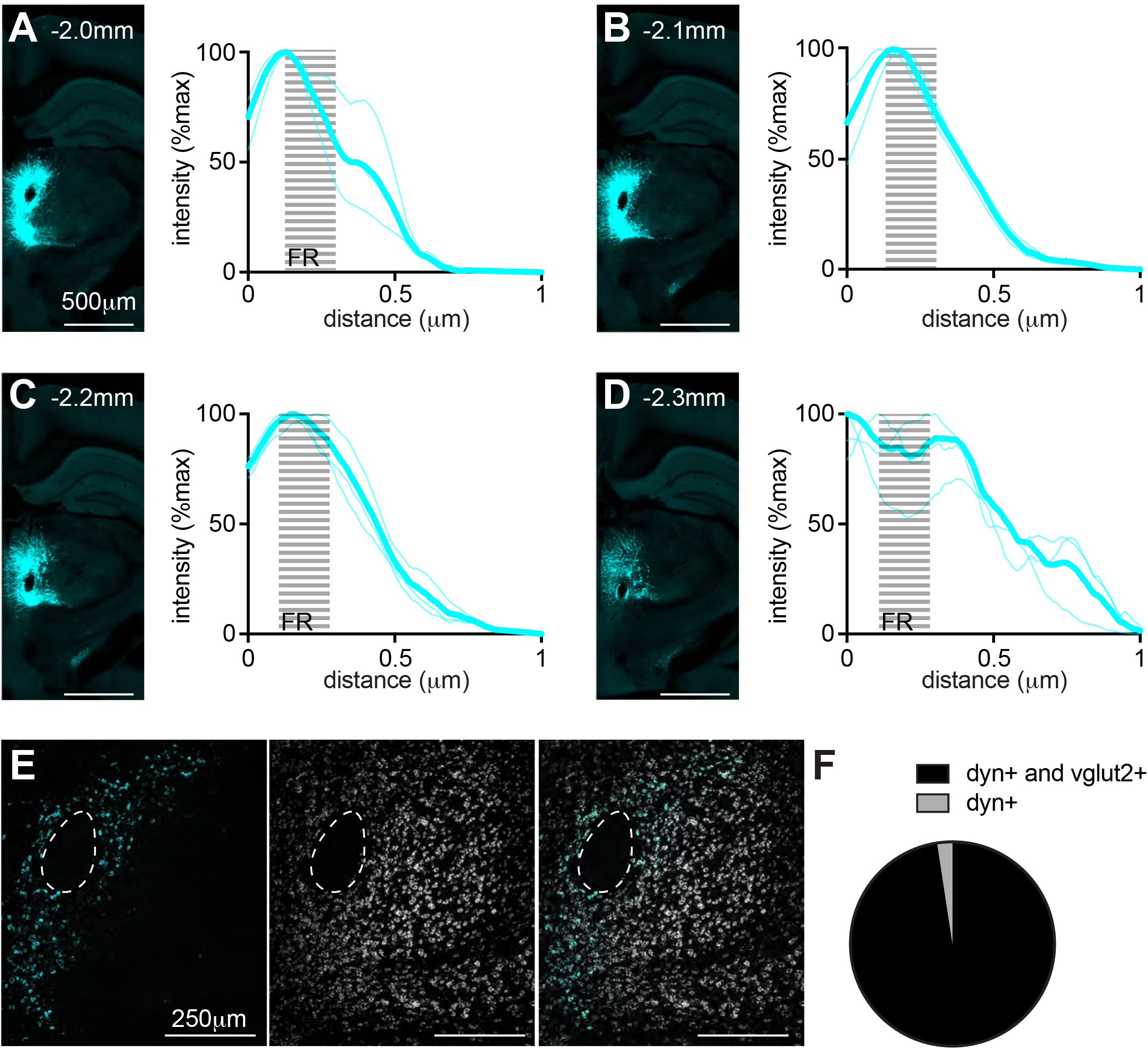
**A-D**, *left*: Example of a coronal section from a *Pdyn-IRES-cre* injected with a creOn-gfp into PF and the quantification of the fluorescent intensity in PF (right) at coronal section -2.0mm (A), -2.1mm (B), -2.2mm (C), -2.3 (D). Thin lines represent peak-normalized data from individual animals and the thick lines show the means for each channel. The dashed grey region represents the FR location (n=3 mice). **E**, Example of ISH for *Pdyn (*cyan), and *Slc17a6 (*white) in the PF of a *Pdyn-IRES-cre* mouse highlighting the overlap of *Pdyn* and vglut2.

**Related to Figure 5:**

**S5:**
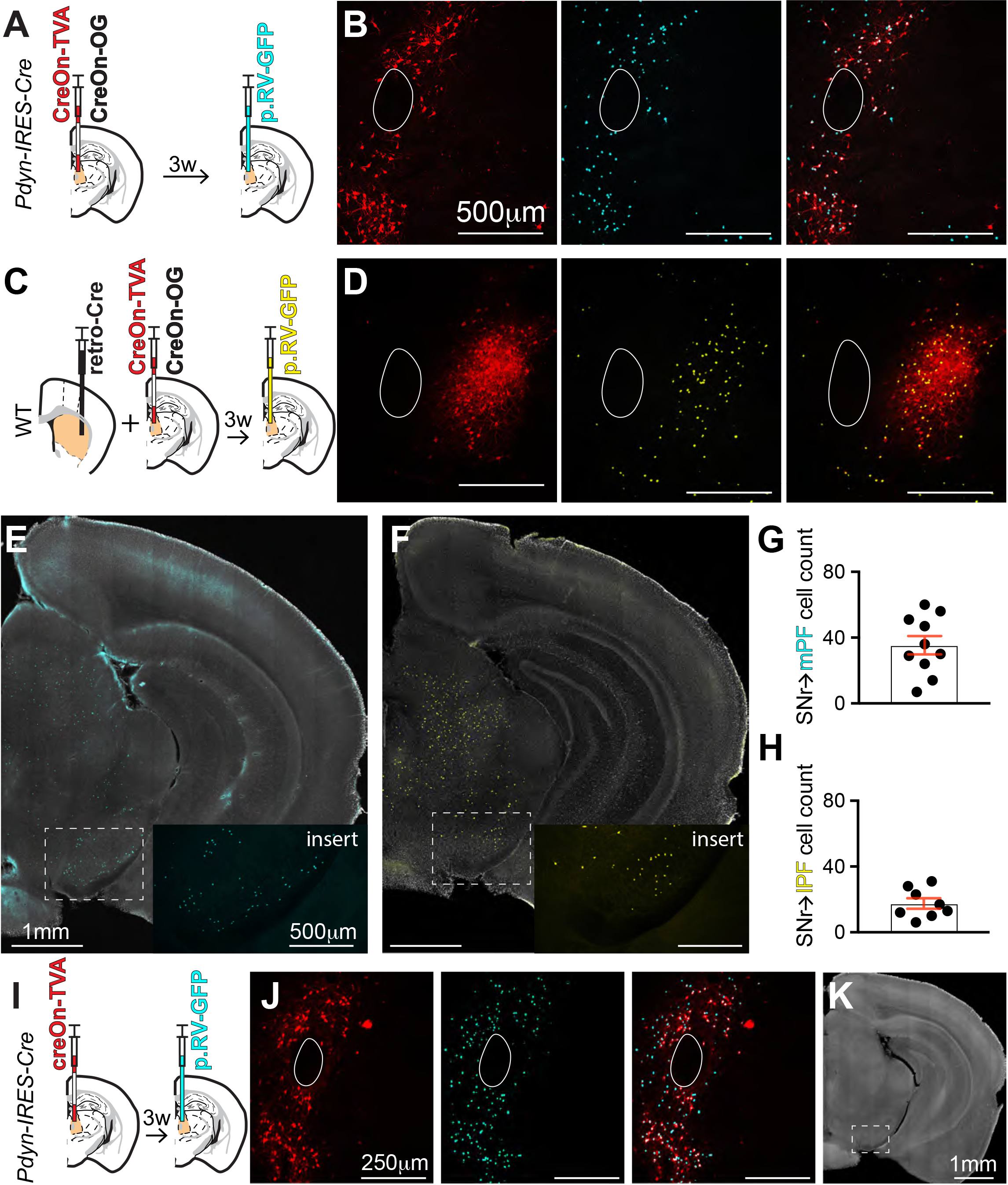
**A**, Experimental design showing a coronal section at -2.1 mm depicting an injection of creOn-TVA and creOn-OG (*left*) followed by an injection 3 weeks later of p.RV-GFP in the PF of a *Pdyn-IRES-cre* mouse (*right*). creOn-TVA, creOn-OG are two helper virus that are necessary for rabies infection and its retrograde transfer, respectively. **B**, *left,* example coronal section at -2.1 mm in PF showing expression of TVA in mPF (red) from the experiment in A. *Middle,* expression of p.RV-GFP (cyan) in lPF. *Right,* overlay of the two channels highlighting that there is no expression of p.RV-GFP in cPF or lPF. (n=3 mice, example shown from one mouse). **C-D**, as in panel (A-B) but expression of creOn-TVA and creOn-OG was induced in lPF using a retrograde traveling AAV with Cre (retro-Cre) injected into dlSTR in a WT mouse. No expression of p.RV-GFP (yellow) was observed in mPF or cPF. (n=3 mice, example shown from one mouse). **E-F**, Coronal sections at -3.7mm from the experiment shown in (A) and (B) respectively highlighting that there is expression of p.RV-GFP+ (cyan or yellow) in the SC and SNr. The SNr is shown in the inset. **G-H**, Number of cells counted in the SNr slices from the experiment shown in (A) (n=354/10/2; p.RV-GFP+ cells/slices/animals) and (B) (n=140/8/2; p.RV-GFP+ cells/slices/animals) highlighting medial SNr➔mPF and lateral SNr➔lPF projections. **I-J**, as in panel (A-B) but with no injection of creOn-OG into a WT mouse. This verifies that the movement of the p.RV-GFP movement upstream and across the synapse is dependent on creOn-OG. **K**, Coronal section at -3.7 mm from the experiment shown in (I) verifying that there is no expression of p.RV-GFP+ outside of PF. SNr is highlighted in a dashed line.

**Related to Figure 6:**

**S6:**
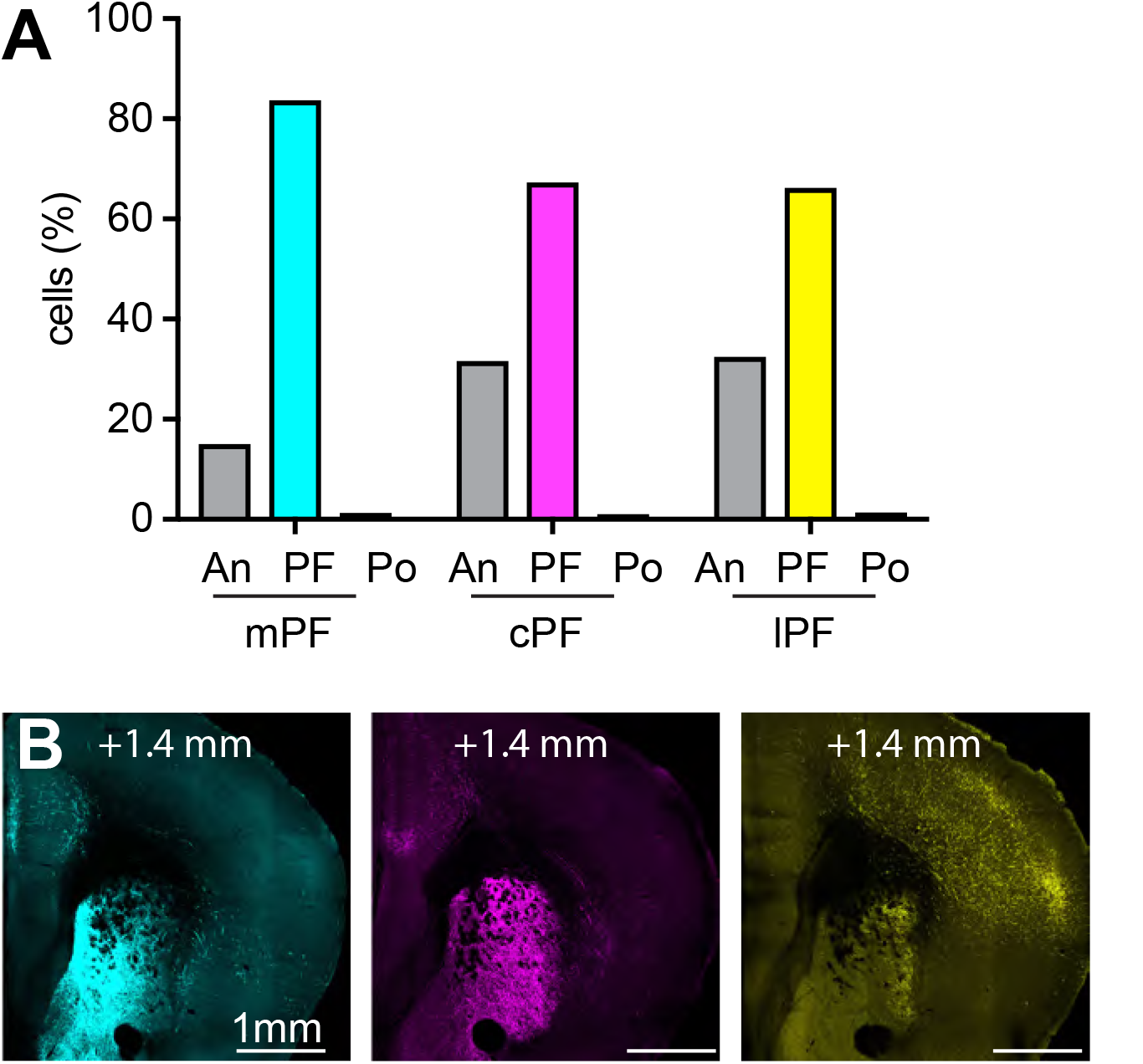
**A**, Percent GFP+ cells found 200 μm anterior to PF (An), in PF, or 100 μm posterior to PF in the animals that were analyzed for their putative inputs from PF to CTX (for mPF=713/1; cPF=1000/1; lPF=2171/1; cells/mice). **B**, Example coronal at +1.4mm from the experimental design and same animals shown in Figure 6 highlighting that the overall topography between PF cell classes and cortex was maintained in between the two regions we analyzed.

**Movie 3:** Example of a whole brain from experimental shown in Figure 6A.

**Related to Figure 7:**

**S7:**
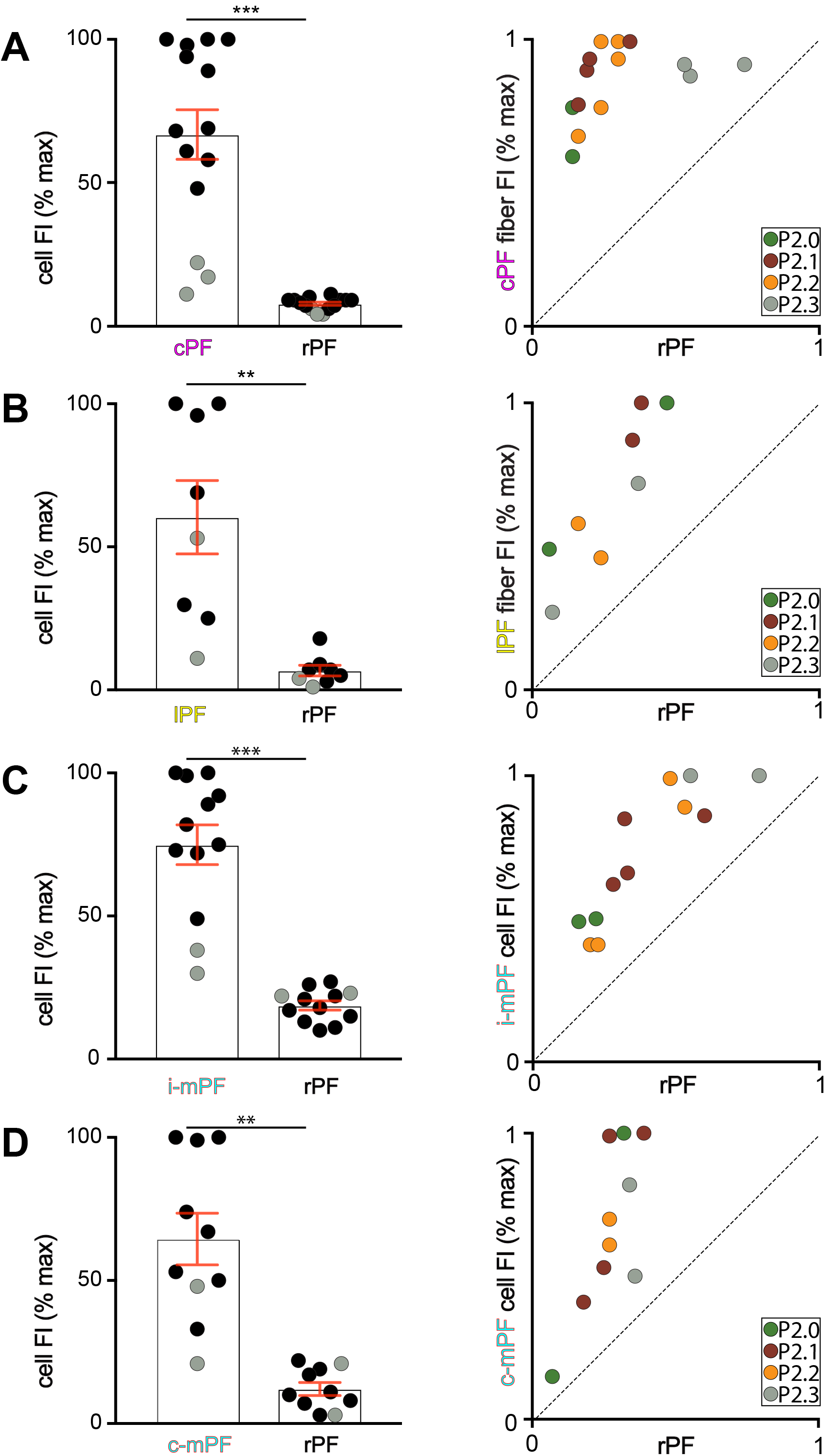
**A**, *left,* Quantification of percentage of the maximal fluorescence intensity of CTB+ in cPF as shown in Figure 7B, compared to that in the rest of PF (rPF). Grey filled circles here (and throughout the figure) represent the analyses of coronal section -2.3mm in PF. P=0.0001; Wilcoxon test. *right,* Quantification of percentage of the maximal fluorescence intensity of GFP labeled axons (Fl) in the cPF ROI, as shown in Figure 7B compared to that in the rest of PF (rPF). Each dot represents one section analyzed with its color reflecting its location within PF. **B-D**, Same as panel (A) but for experiments shown in Figure 7D, Figure 7H, and Figure 7L. For panel B: P= 0.0078, For C: P= 0.0005, For D: P= 0.0020. Wilcoxon test.

**S8:**
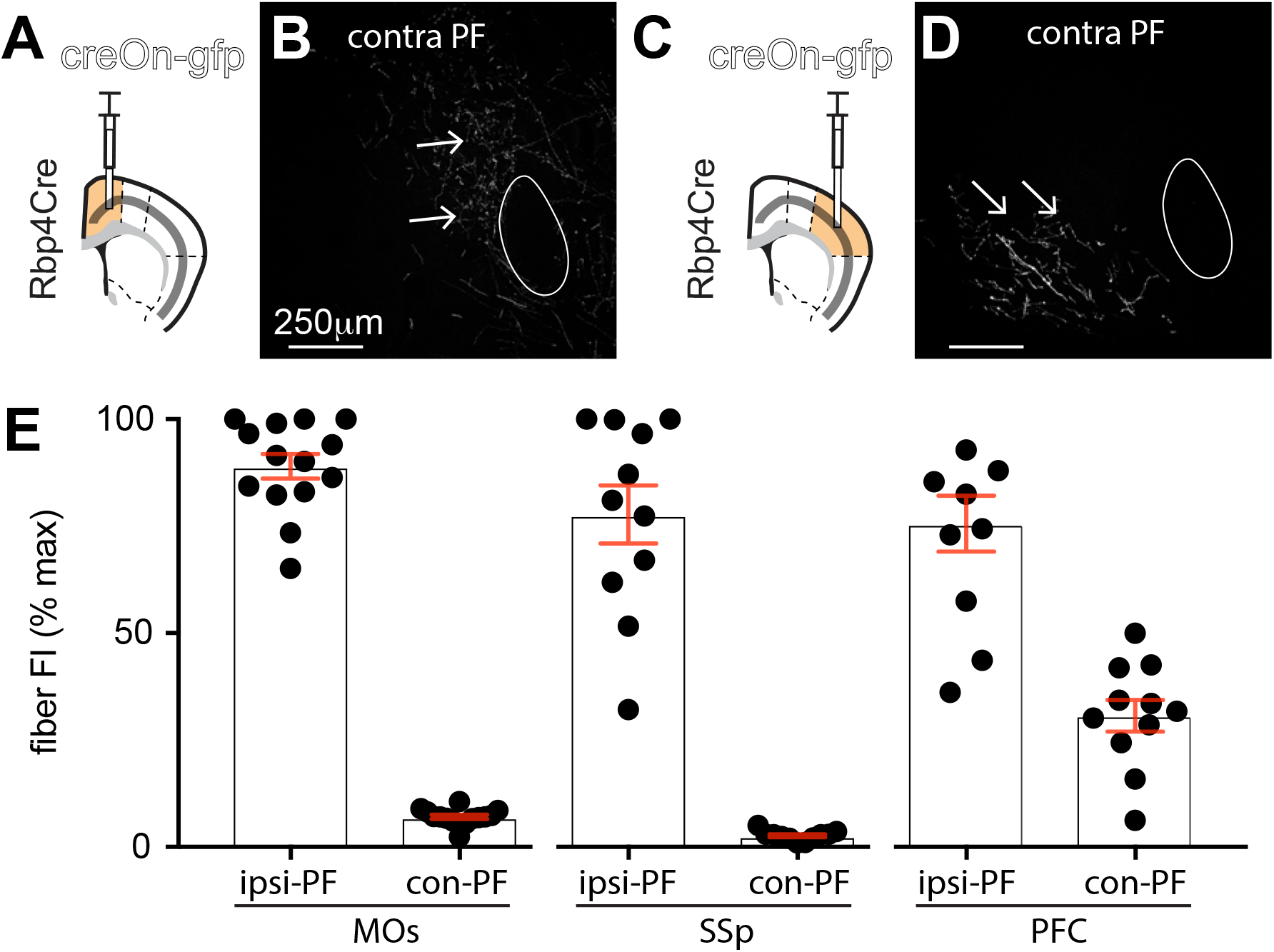
**A**, Experimental design shown of coronal sections at +0.9 mm from a *Rbp4-Cre^+/-^* mouse depicting a viral injection of creOn-GFP (white) into MOs. **B**, Example coronal section from experiment in (A) at -2.1 mm in PF, contralateral to the injection site. The arrows highlight the weak fibers in contralateral PF. **C-D**, As in panel (A) and (B) but for injection of creOn-GFP into SSp. **E**, Quantification of percentage of the maximal FI of GFP labeled axons in PF ipsilateral (IPSI) or PF contralateral (CON) to the injection site in MOs (*left)*, SSp (*middle)*, and PFC (*right)*. This confirmed that MOs➔con-PF and SSp➔con-PF fibers FI are weak compared to PFC➔con-PF fibers.

